# Tumour-derived Ilp8 and Upd3 control intestinal progenitor cells depletion during cachexia in *Drosophila* larvae

**DOI:** 10.1101/2025.04.21.649791

**Authors:** Jennifer Falconi, Miriam Rodríguez-Vázquez, Katrin Strobel, Céline Jahannault-Talignani, Lisa Heron-Milhavet, Patrice Lassus, Charles Géminard, Alexandre Djiane

## Abstract

In animals, tumour development triggers systemic effects, impacting the physiology of distant organs. In *Drosophila* larvae, wing disc neoplastic tumours result in developmental delay and organ wasting reminiscent of cachexia. This paraneoplastic syndrome affects many organs, but its effects on the intestine, a key organ in the regulation of nutrient and energy homeostasis, remain understudied. We describe here that neoplastic tumours also affect the development of the larval midgut, leading to altered cell type numbers, with a depletion of the stem-cell-like Adult Midgut Precursors (AMPs), and a disorganisation of the niche cells which enter precocious differentiation. Importantly, these intestinal cell type alterations are initiated before the onset of reduced food intake, and of muscle and adipose tissue atrophies, and thus represent a new paraneoplastic phenotype. Screening for mediators, we show that tumour derived Ilp8 and Upd3 control AMPs number and niche specification respectively.

## INTRODUCTION

For animals, the development of tumoral tissues represents a specific challenge, where the growing tissues divert nutrient and energy. This could trigger systemic effects where the physiology of the individual and distant organs are affected. Para-neoplastic syndromes are thus adaptations to the tumour presence, and often result in pathological adverse effects.

Cachexia is a systemic metabolic disorder associated with several chronic diseases, including several cancers. It is characterised by an involuntary and rapid weight loss, and is frequently associated with loss of appetite, hyperglycemia, and inflammation. Profound metabolic changes occur during cachexia with the futile usage of energy and macromolecules favouring catabolism. Amongst the most dramatic effects of cachexia are the atrophies of the adipose tissues and of the skeletal muscles (sarcopenia; reviewed in (Baracos *et al*, 2018; Ferrer *et al*, 2023a). In mouse models, mitigating muscle loss by targeting the Activin Receptor IIB (thus blocking the anti-myogenic effect of Myostatin, Zhou *et al*, 2010) or by restoring NAD+ metabolism (Beltrà *et al*, 2023) dramatically prolongs survival of tumour bearing animals. At the organismal level, cachexia involves the complex interplay between the tumours and affected tissues such as muscles and adipose tissue, with also alterations of other organs such as the liver, the brain or the immune system (Baazim *et al*, 2022; Burfeind *et al*, 2020; Xie *et al*, 2022).

Studying cachexia, which implicates intricate exchanges between different organs, requires whole animal models such as mouse models which have identified several potential tumour-derived pro-cachectic factors, and tissue-specific responses (Zhou *et al*, 2010; Ferrer *et al*, 2023b; Liu *et al*, 2024; Kir *et al*, 2014; Sun *et al*, 2024; Xie *et al*, 2022; Bilgic *et al*, 2023); reviewed in (Ferrer *et al*, 2023a). Non-mammalian models of cachexia have also emerged in the genetically tractable *Drosophila melanogaster* characterised by atrophies of the fat body (functional equivalent of the adipose tissue and of the liver), and of the skeletal muscles (Saavedra & Perrimon, 2019). Adult *Drosophila* cachexia models are based either on the graft of RasV12-driven tumorous larval wing discs in the abdomen of adults, or on the generation of tumours in the adult midgut by over-expressing an activated form of Yki in the midgut stem cells (*esg^ts^>YkiS3A*) (Figueroa-Clarevega & Bilder, 2015; Kwon *et al*, 2015). Larval *Drosophila* cachexia models consist of eye discs tumours after expressing the activated Ras (RasV12) in combination with impaired epithelial polarity (*RasV12; scrib-*), or impaired Src signalling on a high sugar diet (*RasV12; dCsk-*) (Lodge *et al*, 2021; Hodgson *et al*, 2021; Newton *et al*, 2020). Adult fly models of cachexia suggested an important role of the Insulin-trapping protein ImpL2 (fly homologue of human IGFBP6&7), of Pvf-1, and of the cytokine Upd3 (IL-6) in mediating fat body and muscle atrophy (Figueroa-Clarevega & Bilder, 2015; Kwon *et al*, 2015; Ding *et al*, 2021; Song *et al*, 2019). In parallel larval models of cachexia supported the role of other tumour secreted factors such as the FGF ligand Bnl (FGF20), and the TGF-β ligand Gbb (BMP6), or of an altered Ecdysone processing (steroid hormone) in mediating larval muscle and fat body atrophy (Newton *et al*, 2020; Lodge *et al*, 2021; Santabárbara-Ruiz & Léopold, 2021), or the relaxin member Ilp8 in promoting anorexia (Yeom *et al*, 2021). These *Drosophila* models have also highlighted important responses in the affected tissues, such as autophagy in the wasted tissues (Khezri *et al*, 2021), metabolic reprogramming in muscles (Saavedra *et al*, 2023), and the contribution of the atrophied fat body to muscle wasting (Bakopoulos *et al*, 2023). Recent reports show that the bloating frequently observed in *Drosophila* cachexia models stems from profoundly affected Malpighian tubules, the functional equivalent kidneys (Xu *et al*, 2024, 2023; Kwok *et al*, 2024), even though kidney dysfunction is not a hallmark of cancer-associated cachexia in patients (Baracos *et al*, 2018; Ferrer *et al*, 2023a).

The intestine, a key metabolic organ implicated in nutrients processing and absorption and in the endocrine control of metabolism, has been shown to be atrophied during cachexia, in mouse and in larval *Drosophila* models (Bindels *et al*, 2018; Holland *et al*, 2022). However, despite a few studies, the impact of cachexia on the intestine remains less studied than the impact on other organs.

The fly digestive system is divided in a foregut, midgut, and hindgut, functionally equivalent to the oesophagus, the intestine, and the colon respectively. Studies performed in the past two decades have established the adult *Drosophila* midgut as an experimental paradigm for the study of stem cells and regenerating epithelia (Ohlstein & Spradling, 2006; Micchelli & Perrimon, 2006; Lemaitre & Miguel-Aliaga, 2013; Miguel-Aliaga *et al*, 2018; Boumard & Bardin, 2021), resulting in a detailed description of the adult midgut biology. In comparison, the larval midgut has been less studied. It consists of a tubular single layered epithelium, surrounded by visceral muscles. From anterior to posterior, five main regions can be distinguished based on anatomical features and changing pH, which might undertake distinct functions (Overend *et al*, 2016). At the cellular level, the larval midgut is composed of four different cell types (Fig. 1A). The large absorptive enterocytes (EC) with large polyploid nuclei (typically 32N), express the marker Pdm1 (also known as Nubbin). The small diploid entero-endocrine cells (EEC) secrete different peptides and hormones, and express the marker Prospero (Pros). The adult midgut precursors (AMPs) represent a pool of diploid cells with stem cell properties set aside during larval life. They originate in the embryo from specific posterior neuroblast-like cells, or endoblasts (Plygawko *et al*, 2025; Micchelli *et al*, 2011), and are scattered as isolated cells along the length of the early larval gut. During the 3^rd^ instar larval stage, AMPs proliferate to generate small islets of 6-7 tightly packed cells surrounded by one Peripheral Cell (PC), which acts as a niche for AMPs preventing their precocious differentiation (Fig. 1B&C; (Mathur *et al*, 2010; Micchelli, 2012). PCs arise from the asymmetric division of AMPs in the islet and both AMPs and PCs express the marker escargot (Esg; (Mathur *et al*, 2010)). During metamorphosis, AMPs across the remodelling gut will proliferate and generate the adult midgut.. Thus, during the short lifespan of the larva, AMPs are normally kept dormant, but can be transiently reactivated to generate new ECs following damage by pathogenic bacterial infection (Houtz *et al*, 2019).

**Figure 1.**
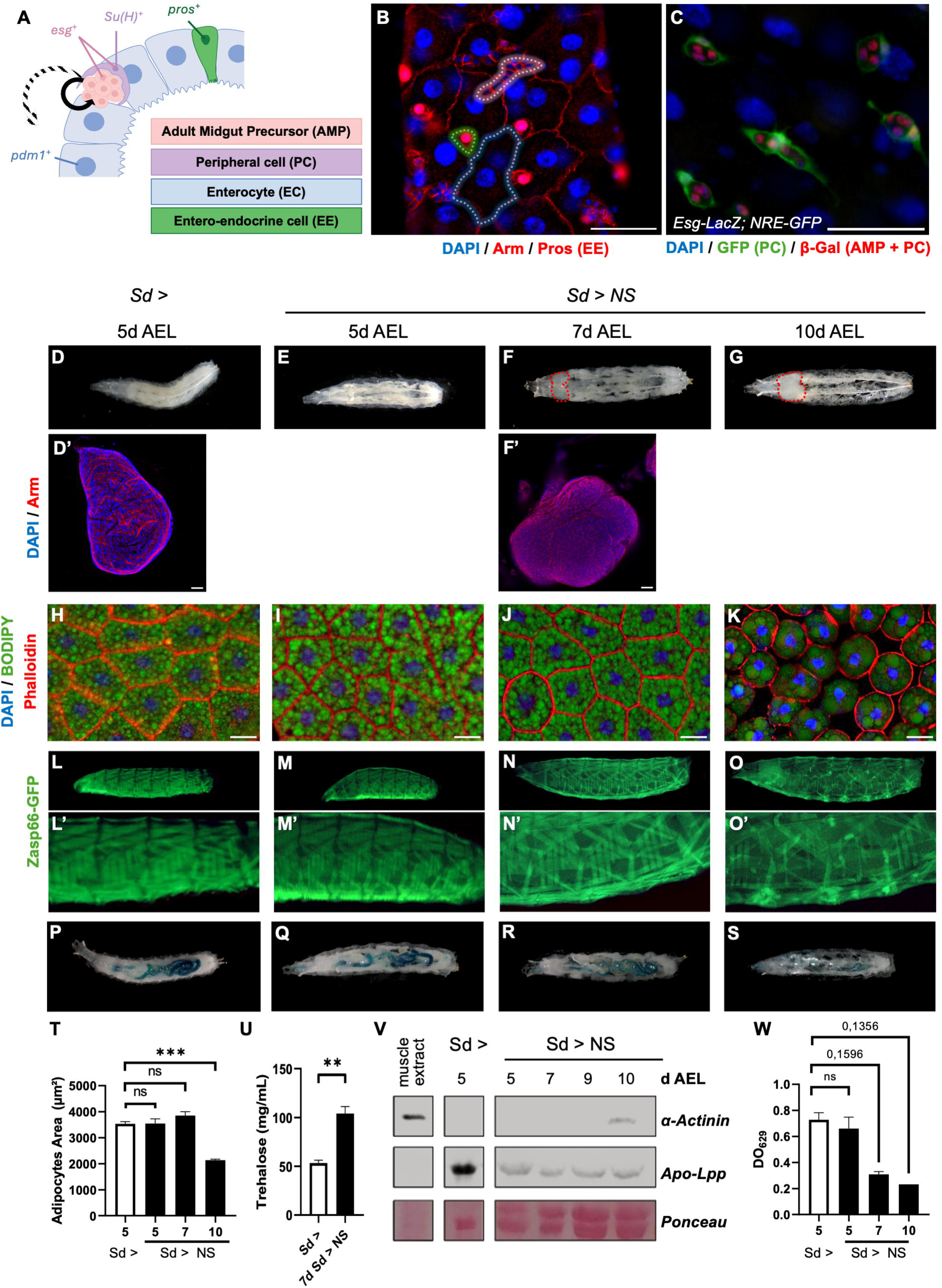
Sd>NS larvae exhibit cachexia-like syndrome. **A.** Schematic of a cross section of the *Drosophila* larval midgut with the different cell types and their associated markers. **B.** Wild-type larval midgut staining at 5d AEL (After Egg Laying) showing nuclei (DAPI, blue), cell cortex (Armadillo, red), and the entero-endocrine cell marker Prospero (red). The doted lines highlight the different cell types: large enterocyte with large nucleus (EC; blue), small entero-endocrine propsero positive cell (EE; green), and cluster of tightly packed adult midgut precursor small cells (AMP; red). White bar: 50 µm. **C.** Wild-type Larval midgut staining at 5d AEL showing nuclei (DAPI, blue), the NRE-GFP positive peripheral cells (PC; green), and the expression of the AMP and PC marker Esg (Esg-LacZ; red). White bar: 50 µm. **D-G.** Timeline of larvae morphology in response to neoplastic wing disc tumours with activated Notch and mutant for *scribble* (Sd>NS) at 5d (E), 7d (F), and 10d AEL (G) compared to wild type 5d AEL Sd> controls (D). (D-G) Whole larva, head on the left. Red dotted lines in F&G indicate the overgrowing wing disc tissue. (D’, F’) Corresponding dissected wing imaginal discs and stained for Armadillo (red), and DAPI (nuclei, blue). White bar: 50 µm. **H-K.** Staining of larval fat body showing Actin (red), lipid droplets (BODIPY, green), and nuclei (DAPI, blue). Control Sd> larvae at 5d AEL (H), and cachectic Sd>NS larvae at 5d AEL (I), 7d AEL (J) and 10d AEL (K). White bar: 50 µm. **L-O.** Larval body wall muscles monitored using the Zasp66:GFP reporter in live larvae (green). White bar: 200 µm. (L’-O’) higher magnification of images shown in (L-O). **P-S**. Food intake monitored with blue food colouring. **T.** Quantification of adipocyte cell size from images shown in (H-K). Biological replicates n ≥ 5. Error-bars show standard error of the mean (sem). Normality was tested with a Shapiro-Wilk Test. One-way ANOVA statistical test. *** p < 0.001, ns = not significant. **U.** Quantification of trehalose concentration in the haemolymph. Biological replicates n = 3. Error-bars show standard error of the mean (sem). Normality was tested with a Shapiro-Wilk Test. Unpaired t-test. ** p < 0.01. **V.** Western blot of whole protein extracts of hemolymph (5µl samples) from Sd> control larvae at 5d AEL, and Sd>NS cachectic larvae at 5, 7, 9, and 10d AEL, and monitoring the presence of the muscle protein α-Actinin. Apo-Lpp, and Ponceau are used as loading control. **W**. Quantification of blue food from larvae shown in (P-S). Each measure represents a pulled sample of 8 larvae. Error-bars show standard error of the mean (sem). Normality could not be verified with a Shapiro-Wilk Test. Kruskal-Wallis with exact p-value plotted.

Here we describe a new larval model of cachexia, based on Notch-based tumoral overgrowth of imaginal wing discs (Logeay *et al*, 2022). We show that all the major hallmarks of cachexia described in the Ras-based models in the eye are also found in our models, demonstrating that the para-neoplastic syndromes observed are not oncogene, nor eye disc specific. During cachexia, we describe a profound alteration of the midgut cellular composition with abnormal PCs and fewer ECs which enter differentiation. Molecularly, we show that changes in PCs and ECs are driven by Ilp8 and Upd3 produced by the tumours. Our studies thus show that tumours, through the production of Ilp8 prevent AMP proliferation and through the pro-inflammatory cytokine Upd3 promote PC inappropriate engagement and depletion.

## RESULTS

### Larval wing disc overgrowth induce wasting and cachexia-like syndrome

We previously described a larval wing disc tumour model based on the activation of the Notch pathway (UbxFlp-MARCM model; (Logeay *et al*, 2022)). The induction of wing discs clones overexpressing an activated form of the Notch receptor (Notch intra cellular domain, Nicd) combined with a loss of function allele for *scrib*, resulted in neoplastic overgrowing discs (UbxFlp-MARCM with UAS Nicd and FRT82B *scrib1*; Ubx>NS discs; Supplemental Fig. S1A-D) (Logeay *et al*, 2022). Interestingly, animals bearing Ubx>NS tumours failed to pupate and had an extended larval life of up to 20 days after egg laying (AEL) instead of the normal 5 days. Ubx>NS animals presented several phenotypes reminiscent to cachexia. First, at 10d AEL, the fat body in Ubx>NS larvae was severely atrophied and more transparent with smaller and less cohesive (rounder cells) adipocytes. Adipocytes were slightly smaller and adopted a round shape smaller in size (≈ 20% smaller; Supplemental Fig. S1E-H). The corresponding larvae had lower total glycogen content and increased circulating trehalose concentration (Supplemental Fig. S1I&J). Strikingly, starting around 10d AEL, muscle proteins such as α-Actinin or Myosin were detected in the haemolymph by western blot analysis, a sign of muscle damage similarly to what we reported recently for fat body stress induced muscle damage (Supplemental Fig. S1K; (Rodríguez-Vázquez *et al*, 2024)). The fat body and muscle atrophies observed in Ubx>NS animals were progressive phenotypes and fat body atrophy appeared slightly earlier than muscle atrophy, consistent with a recent report suggesting that the atrophied fat body fails to provide the extracellular matrix component Viking (Collagen IV) important for muscle integrity in *Drosophila* larvae (Bakopoulos *et al*, 2023). Finally, even though not related to current pathological symptoms of cachexia in patients, Ubx>NS animals were also heavily bloated, a phenotype that has been recently associated with nephrocytes dysfunction either as a consequence of tumour derived anti-diuretic secreted peptides, or nephron stem cell abnormal differentiation in adult *Drosophila* cachexia models (Xu *et al*, 2024, 2023; Kwok *et al*, 2024), and which was reported in other cachexia larval models (Santabárbara-Ruiz & Léopold, 2021).

However, the complex genetics of Ubx>NS make these animals complicated to use for functional manipulations. We thus replicated the Ubx>NS conditions using the Scalloped-Gal4 strong wing disc pouch driver (Sd-Gal4). Sd-Gal4 driving UAS Nicd and a UAS RNAi targeting *scribble* (Sd>NS), resulted in neoplastic growth, with the animals presenting all the cachexia phenotypes presented for the Ubx>NS model: extended larval life and bloating (Fig. 1D-G), smaller and rounder adipocytes (Fig. 1H-K&T), muscle atrophy and disorganisation (as evidenced with the live muscle marker Zasp66-GFP) with release of α-Actinin in the haemolymph (Fig. 1L-O&V), and hypertrehalosemia (Fig. 1U). We also monitored food intake in Sd>NS larvae using blue-dyed food which showed that Sd>NS larvae started to eat less than controls around 7d AEL. Importantly there was no difference in feeding amount at 5d AEL between Sd>NS and Sd controls (Fig. 1P-S&W).

These different observations suggest that *Drosophila* larvae with wing discs NS neoplastic tumours suffer from cachexia-like syndrome (Ferrer *et al*, 2023a; Saavedra & Perrimon, 2019; Baracos *et al*, 2018). This new Notch-driven wing disc model thus extends the observations made with the Ras-driven eye disc tumours (Lodge *et al*, 2021; Santabárbara-Ruiz & Léopold, 2021; Hodgson *et al*, 2021; Yeom *et al*, 2021; Khezri *et al*, 2021; Bakopoulos *et al*, 2023; Newton *et al*, 2020), thus validating that the observations made with larval cachexia models are not disc or driving oncogene specific, even though some differences might exist in the intricate molecular mechanisms leading to cachexia.

We decided here to focus our analysis on the *Drosophila* larval gut, an organ that has been relatively less studied than other organs during the establishment of cachexia-like wasting syndromes or para-neoplastic syndromes in *Drosophila* models. Importantly, we could not detect any activity of our Ubx-MARCM or Sd-Gal4 drivers in the larval midgut (Supplemental Fig. S2A-D), confirming that any potential phenotype observed in the larval midgut in response to NS tumours was not due to leakiness.

### Cachectic midguts contain fewer *escargot* positive cells

We thus investigated the cellular composition of the cachectic larval midgut. We probed the relative proportions of midgut cell populations by performing qRT-PCR using cell type specific genes: *esg* specific of the AMPs and of the peripheral niche cell (PCs), *pdm1* specific of the enterocytes, and *pros* specific of the endocrine cells (Fig. 1A). In Sd>NS midguts at 5d AEL and at 7d AEL, we observed a strong reduction of the relative expression of *esg*, and an increase in the expression of *pdm1,* when compared to Sd-Gal4 control midguts at 5d AEL. The expression of *pros* remained unchanged (Fig. 2A). This suggests that either there were fewer AMPs and PCs *esg* positive cells, or that the level of expression of *esg* in AMPs and PCs was reduced (or both). We then evaluated AMPs number by immunostaining using an Esg-LacZ reporter line focusing on the anterior midgut (anterior to the acidic copper cell region). Consistent with this reduced *esg* expression, Sd>NS midguts had fewer AMPs at 5d and 7d AEL than Sd-Gal4 controls (5d AEL; Fig. 2B-D). At 7d AEL, NS midguts had a decreased density of islet and a decreased number of cells per islet (Fig. 2G&H). Similar phenotypes were also seen with the Ubx>NS model in which we also observed a reduction of *esg* expression by qPCR at 7d AEL (Supplemental Fig. S3A). In the Ubx-MARCM model, we evaluated AMP numbers, by counting the number of cells with a small nucleus and present in islets with high Armadillo staining (AMPs surrounded by PC with strong adherens junctions). Indeed cells with a large nucleus are Enterocytes, and isolated cells with small nucleus positive for Prospero staining are the Entero-Endocrine cells (Supplemental Fig. S3B&C). Quantifying islets and AMP numbers, we observed roughly similar number of islets but fewer AMP cells with small nuclei per islets, thus resulting in reduced total AMP numbers consistent with the decreased *esg* expression (Supplemental Fig. S3D-F). These results demonstrate that the effect observed on the cellular composition of the cachectic guts were not due to unspecific effects of the Sd-Gal4 driver and were the consequence of the wing discs tumours acting remotely on the midgut. Altogether these results suggest that neoplastic tumours induce a reduction in AMP numbers at 5d AEL, a defect that is not corrected later. This early reduction in AMPs could be interpreted as a delay in AMP proliferation within islets. Indeed, we observed that midguts from NS animals had a lower expression level of the mitosis key regulator *polo* (Fig. 2A), suggesting that the midgut of NS animals contains fewer cells able to proliferate and undergo mitosis.

**Figure 2.**
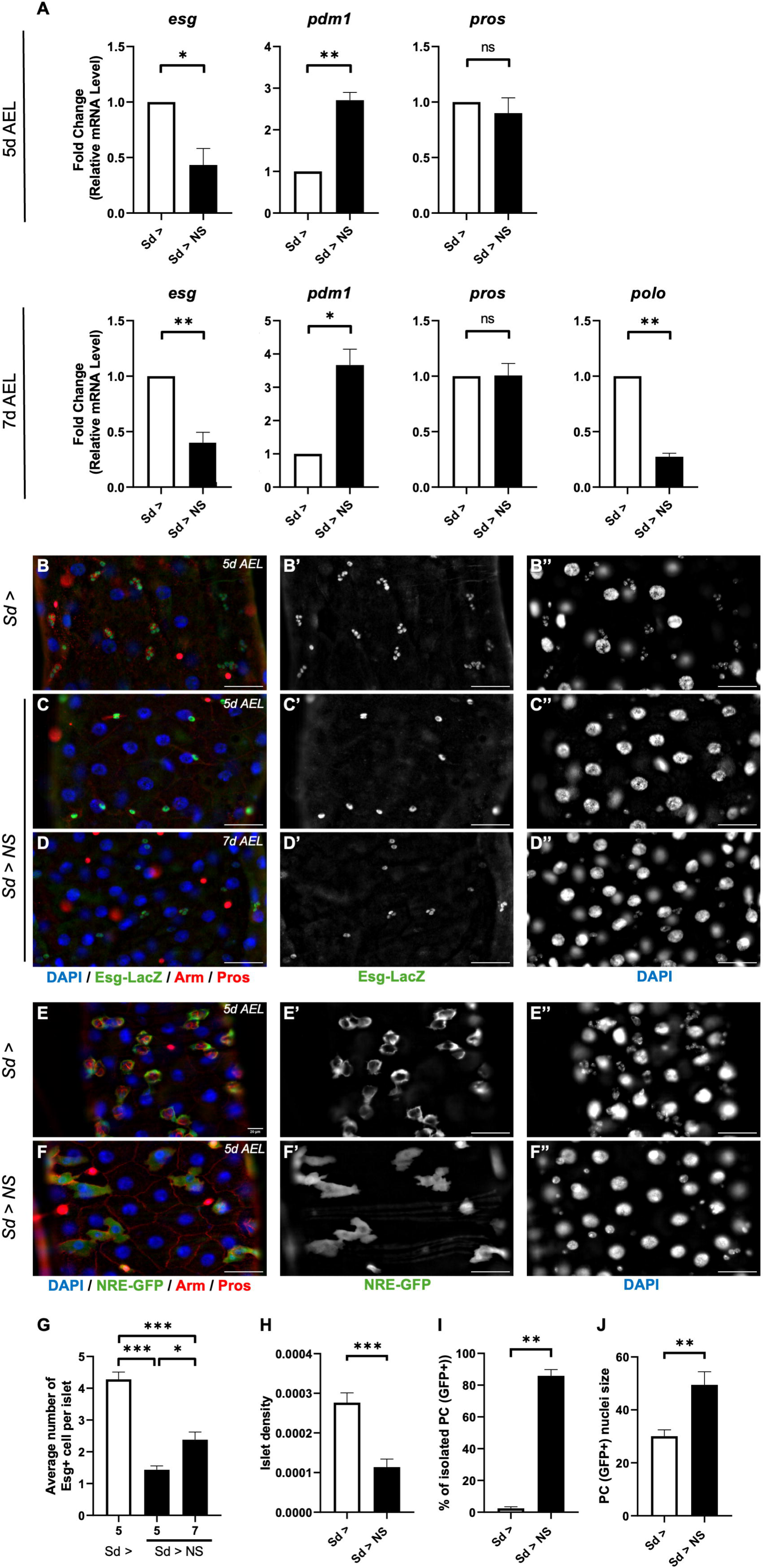
Larval midgut cell type alterations during cachexia. **A.** Expression of the cell type markers *esg*, *pdm1*, *pros*, and of the cell cycle marker *polo* in larval midguts at 5d AEL (upper panels) and 7d AEL (lower panels) monitored by qRT-PCR. Reference gene for normalization: *rp49/RpL32*. Biological replicates n ≥ 3. Error-bars show standard error of the mean (sem). Normality was tested with a Shapiro-Wilk Test. One-sample t-test test. ** p < 0.01, * p < 0.05, ns = not significant. **B-D.** Larval midgut staining showing the EE marker Pros (red), the AMP&PC marker Esg (Esg-LacZ, green in B-D, white in B’-D’), and nuclei (DAPI, blue in B-D, white in B’’-D’’). Membranes are marked with Arm (red). Control Sd-Gal4 midguts at 5d AEL (B), and cachectic Sd>NS midguts at 5d AEL (C) and 7d AEL (D). White bar: 50 µm. **E-F.** Larval midgut staining showing the EE marker Pros (red), the PC marker NRE-GFP (green in E-F, white in E’-F’), and nuclei (DAPI, blue in E-F, white in E’’-F’’). Membranes are marked with Arm (red). Control Sd-Gal4 midguts (E) and cachectic Sd>NS midguts (F) at 5d AEL. White bar: 50 µm. **G-H.** Quantification of number of Esg positive cells per islet (G) and of islet number per area unit (density; H) from images shown in (B-D). Biological replicates n ≥5. Error-bars show standard error of the mean (sem). Normality was tested with a Shapiro-Wilk Test. One-way ANOVA statistical test (G) and T-test statistical test (H). *** p < 0.001, * p < 0.05. **I-J.** Quantification of PC isolation frequency (I) and PC nuclei size expressed in arbitrary pixel number (J) from images shown in (E-F). Biological replicates n ≥5. Error-bars show standard error of the mean (sem). Normality was tested with a Shapiro-Wilk Test. Mann Whitney test (I) and T-Test statistical test (J). ** p < 0.01.

We then investigated the niche represented by the PCs in the midgut of cachectic animals. PCs are marked by the specific expression of the Notch activity reporter *NRE-GFP*. In normal midguts, *NRE-GFP* positive PCs consisted of large cells engulfing several AMPs (Fig. 2E). However, at 5d AEL in Sd>NS midguts, *NRE-GFP* positive PCs appeared mishappen with a straighter shape that no longer encircled AMPs (Fig. 2F&J), suggesting that niche failure could be one cause of this reduction in AMPs *esg* positive cells, and the inability of NS animals to restore the normal set of AMPs per islets at later stages. Importantly, these phenotypes of mishappen PCs and fewer AMPs, were detected at 5d AEL, earlier than the onset of anorexia in the Sd>NS or Ubx>NS models (Fig. 1P-S&W), suggesting that this cell population remodelling of the larval midgut during cachexia is not a consequence of reduced food intake.

### PCs engage in precocious differentiation during cachexia in *Drosophila* larvae

The low *esg* expression in cachectic NS midguts could thus be explained by a lower number of AMPs which fail to proliferate. The proliferation defect rather than increased cell death is supported by the low *polo* expression (Fig. 2A), and by the fact that we could not detect any sign of increased cell death in the cachectic guts, without specific signal for caspase activation (Dcp-1 staining; Supplemental Fig. S3G&H).

But the low *esg* expression could also be a consequence of a depletion in PCs, the other cell type expressing *esg* in the larval midgut, and our studies support their precocious differentiation. Indeed, the PCs had bigger nuclei in NS larval midguts compared to controls, showing that these PCs were starting to become polyploid, strongly suggesting early commitment towards the enterocyte fate (Fig. 2J), consistent with the relative increased expression of the enterocyte marker pdm1 (Fig. 2A). We next monitored *esg* expression in these misshapen PCs using the *Esg-LacZ* reporter expecting to observe a decrease. While the β-galactosidase staining was indeed less intense in the nuclei of NS PCs (signal less concentrated; Supplemental Fig. S4A-C), when integrating the increased size of the nuclei (Supplemental Fig. S4D), the total quantity of β-galactosidase staining was not changed in NS PCs compared to controls (Supplemental Fig. S4E). However, it should be noted that the β-galactosidase protein monitored here is relatively stable and is likely not suited to follow rapid changes in *esg* expression.

### Tumour-derived Ilp8 mediates low AMP numbers

We then searched for the signals responsible for the low AMP numbers during cachexia. Mining the transcriptomes of the Ubx>NS tumours (Logeay *et al*, 2022) identified several genes coding for secreted factors specifically up-regulated in cachectic tumours compared to control discs (Fig. 3A): *bnl, spi, ITP, ImpL2, pvf1, upd1, upd2, upd3,* and *Ilp8*. Several of these secreted molecules such as Bnl, ImpL2, Pvf1, and Upd3 have been previously implicated in mediating some of the cachectic effects of growing tumours in larval (Bnl; (Newton *et al*, 2020)) and adult cachexia models (ImpL2, Pvf1, Upd3; (Figueroa-Clarevega & Bilder, 2015; Kwon *et al*, 2015; Ding *et al*, 2021; Song *et al*, 2019). ITP, Pvf1, and Upd3 were also described to mediate Malpighian tubules dysfunction and bloating (Kwok *et al*, 2024; Xu *et al*, 2024, 2023). However, other ones such as Spi have not been studied as potential tumour-derived pro-cachectic factor. We thus invalidated each of these factors by RNAi in the Sd>NS model and screening by qPCR for a reversal of the *esg* down-regulation in the larval gut that was observed in cachectic midguts (see Fig. 2A; we validated that these factors were also up-regulated in Sd-NS discs; Supplemental Fig. S5A). Amongst the different factors screened, we observed a partial reversal of the phenotype after knocking-down *upd3*, and *Ilp8* (Fig. 3B&C and Fig. 4A).

**Figure 3.**
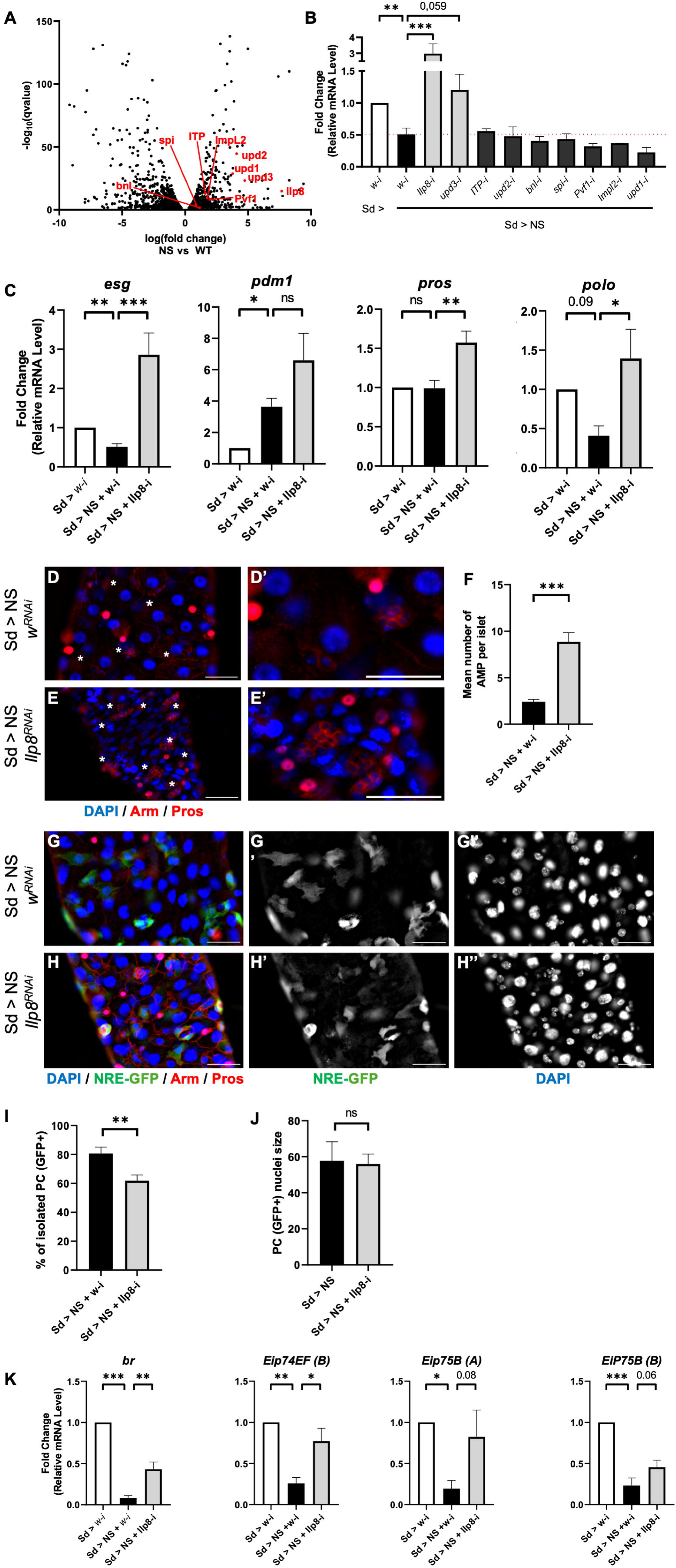
Tumour-derived Ilp8 restricts AMP numbers in cachectic midguts. **A.** Volcano plot showing differentially expressed genes between Ubx>NS and Ctrl wing discs determined by DE-Seq2 from the dataset GEO GSE18533, and filtered for up-regulated secreted ligands shown in red. **B.** Monitoring of *esg* expression in larval midguts at 7d AEL after RNAi-mediated knock down of the different secreted proteins in NS wing disc tumours. Biological replicates n ≥ 2. Error-bars show standard error of the mean (sem). Normality could not be verified with a Shapiro-Wilk Test. Kruskal-Wallis statistical test. ** p < 0.01, *** p < 0.001. **C.** Expression of the cell type markers *esg*, *pdm1*, *pros*, and of the cell cycle marker *polo* in larval midguts at 7d AEL after *Ilp8* RNAi in wing discs and monitored by qRT-PCR. Reference gene for normalization: *rp49/RpL32*. Biological replicates n ≥ 5. Error-bars show standard error of the mean (sem). One-way ANOVA statistical test. *** p < 0.001, ** p < 0.01, * p < 0.05, ns = not significant. **D-E.** Staining of larval midguts at 7d AEL showing the EE marker Pros (red), the membrane marker Armadillo (red), and nuclei (DAPI, blue). Cachectic Sd>NS midguts with a control *white* RNAi (D) or *Ilp8* RNAi (E). White stars highlight AMP islets; white bar: 50 µm. (D’-E’) higher magnification of images shown in (D-E); white bar: 50 µm. **F.** Quantification of number of AMP cells with small nuclei per islet from images shown in (D-E). Biological replicates n ≥ 5. Error-bars show standard error of the mean (sem). Normality was tested with a Shapiro-Wilk Test. Unpaired T-test statistical test. *** p < 0.001. **G-H.** Staining of larval midguts at 5d AEL showing the EE marker pros (red), the membrane marker Armadillo (red), the PC marker NRE-GFP (green in G-H, white in G’-H’), and nuclei (DAPI, blue in G-H, white in G’’-H’’). Cachectic Sd>NS midguts with a control *white* RNAi (G) or *Ilp8* RNAi (H). White bar: 50 µm. **I-J.** Quantification of PC isolation frequency (I) and PC nuclei size expressed in arbitrary pixel number (J) from images shown in (G-H). Biological replicates n ≥ 5. Error-bars show standard error of the mean (sem). Normality was tested with a Shapiro-Wilk Test. Mann Whitney statistical test. ** p < 0.01 ns = not significant. **K.** Expression of the ecdysone signalling targets *broad*, *Eip74*, and *Eip75* in the midguts at 7d AEL of Sd>NS cachectic *Drosophila* larvae with a control *white* RNAi or *Ilp8* RNAi, and monitored by qRT-PCR. Reference gene for normalization: *rp49/RpL32*. Biological replicates n = 4. Error-bars show standard error of the mean (sem). Normality was tested with a Shapiro-Wilk Test. One-way ANOVA statistical test. *** p < 0.001, ** p < 0.01, * p < 0.05.

**Figure 4.**
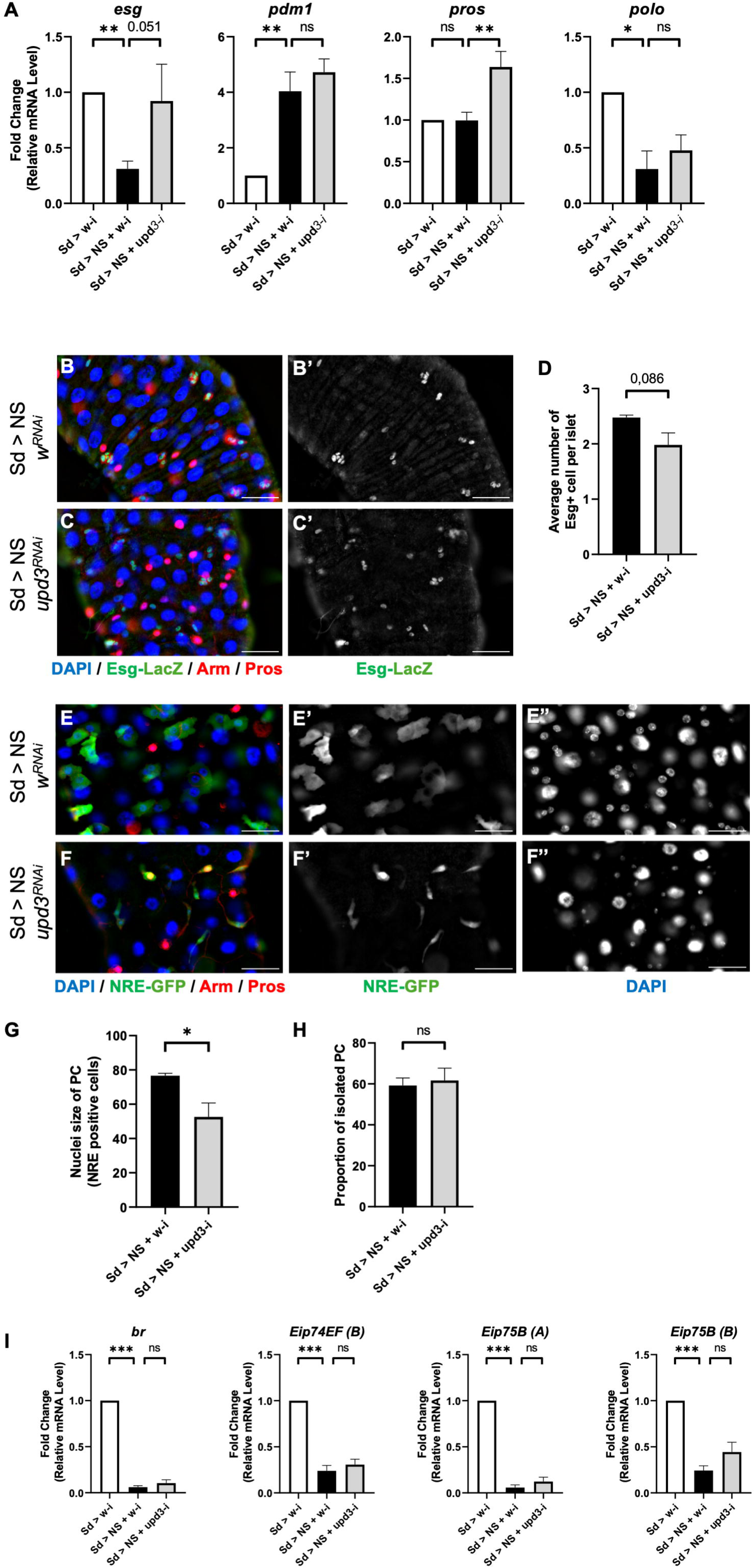
Tumour-derived Upd3 promotes PC differentiation. **A.** Expression of the cell type markers *esg*, *pdm1*, *pros*, and of the cell cycle marker *polo* in larval midguts at 7d AEL after *upd3* RNAi in wing discs and monitored by qRT-PCR. Reference gene for normalization: *rp49/RpL32*. Biological replicates n = 5. Error-bars show standard error of the mean (sem). Normality was tested with a Shapiro-Wilk Test. One-way ANOVA statistical test. ** p < 0.01, * p < 0.05, ns = not significant. **B-C.** Staining of larval midguts at 7d AEL showing the EE marker Pros (red), the membrane marker Armadillo (red), the AMP and PC marker Esg (Esg-LacZ, green in B-C, white in B’-C’), and nuclei (DAPI, blue). Cachectic Sd>NS midguts with a control *white* RNAi (B) or *upd3* RNAi (C). White bar: 50 µm. **D.** Quantification of number of Esg positive cells per islet from images shown in (B-C). Biological replicates n ≥ 5. Error-bars show standard error of the mean (sem). Normality was tested with a Shapiro-Wilk Test. Unpaired t test with Welch’s correction, exact p-value plotted. **E-F.** Staining of larval midguts at 5d AEL showing the EE marker Pros (red), the membrane marker Armadillo (red), the PC marker NRE-GFP (green in E-F, white in E’-F’), and nuclei (DAPI, blue in E-F, white in E’’-F’’). Cachectic Sd>NS midguts with a control *white* RNAi (E) or *upd3* RNAi (F). White bar: 50 µm. **G-H.** Quantification of PC nuclei size expressed in arbitrary pixel number (G) and PC isolation frequency (H) from images shown in (E-F). Biological replicates n = 3. Error-bars show standard error of the mean (sem). Normality was tested with a Shapiro-Wilk Test. Mann Whitney (G) and Unpaired t test (H). * p < 0.05, ns = not significant. **I.** Expression of the ecdysone signalling targets *broad*, *Eip74*, and *Eip75* in the midguts at 7d AEL of Sd>NS cachectic *Drosophila* larvae with a control *white* RNAi or *upd3* RNAi, and monitored by qRT-PCR. Reference gene for normalization: *rp49/RpL32*. Biological replicates n = 4. Error-bars show standard error of the mean (sem). Normality was tested with a Shapiro-Wilk Test. One-way ANOVA statistical test. *** p < 0.001, ns = not significant.

The RNAi-mediated knock-down of *Ilp8* in the NS wing discs tumours rescued efficiently the numbers of AMPs per clusters at 7d AEL as monitored by the number of clustered cells with small nuclei (Fig. 3D-F). Accordingly, *Ilp8* knock-down rescued polo expression consistent with a rescue in the number of proliferating AMP cells (Fig. 3C). Conversely the disorganization of the niche and the PCs detachment were only moderately suppressed by *Ilp8* RNAi at 5d AEL (monitored by NRE-GFP labelling of PCs; Fig. 3G-I). Importantly, *Ilp8* RNAi did not rescue the increase in nuclear size of the misshapen PCs in Sd>NS midguts (Fig. 3J). Together, these results suggest that Ilp8 prevents AMP expansion, but is not involved in PCs detachment and engagement towards EC differentiation. Importantly, removing *Ilp8* in NS tumours did not alter their growth (Supplemental Fig. S5B-D), demonstrating that the effects observed on the midgut were not caused by smaller tumours.

Increased expression of Ilp8 in overgrowing or damaged imaginal discs control developmental timing by activating remotely Lgr3+ positive neurons in the central brain which in turn prevent the secretion of ecdysone by ring gland cells (Colombani *et al*, 2015, 2012; Garelli *et al*, 2012, 2015; Vallejo *et al*, 2015). Interestingly, ecdysone signalling during larval stages is required for AMP proliferation and to reach the correct number of AMPs per islet (Micchelli *et al*, 2011). We thus favour a model where the low AMP numbers observed in animals bearing NS tumours at 5d AEL is the consequence, at least in part, of lower ecdysone signalling following developmental delay induced by Ilp8. Accordingly, we observed a decreased expression of several ecdysone signalling transcriptional targets such as *broad*, *Eip74, or Eip75*, in NS guts at 7d AEL compared to WT controls, which was rescued by *Ilp8-RNAi* (Fig. 3K).

### Tumour-derived Upd3 mediates PC differentiation

In the targeted screen for NS tumours secreted factors controlling *esg* expression in larval midguts, we also identified *upd3* (Fig. 4A). We thus monitored the morphology of the AMPs islets upon *upd3-RNAi* in the tumours. Surprisingly, we did not observe any significant rescue in the number of AMPs per islets, indicating that Upd3 produced by the tumours was not mediating the reduction in AMPs number (Fig. 4B-D). Consistently, we did not observe any rescue to polo expression (Fig. 4A), further demonstrating that Upd3 did not control AMP proliferation. Furthermore, the reduction in the expression of Ecdysone signalling targets was not affected by *upd3* knock-down, further suggesting that in this system, Upd3 does not act on developmental timing and AMPs proliferation (Fig. 4I; (Romão *et al*, 2021)). It is noteworthy that using RNAi we only achieved a partial knock-down of *upd3* which levels remained more elevated than baseline levels of WT controls (Supplemental Fig. S5E’). Focusing on PCs, *upd3* knock-down in NS tumours prevented the increase in PCs nuclear size, suggesting that NS tumours-derived Upd3 triggered PC polyploidisation and engagement towards an early EC fate. (Fig. 4E-G). These different results suggest that Upd3 promotes low relative *esg* expression in midguts, either through the loss of *esg* expression or through the relative increase in *pdm1* expression in PCs following their early differentiation. It is noteworthy that removing *upd3* did not have any significant effect on PC isolation, and PCs remained elongated failing to wrap around the AMP clusters (Fig. 4E,F,H). Whether PC isolation is caused by another signal yet to be identified, or whether the *upd3* levels left after the partial knock-down were sufficient, remains to be explored.

Importantly, removing *upd3* in NS tumours did not alter their growth (Supplemental Fig. 5B,C,E), demonstrating that the effects observed on the midgut were not caused by smaller tumours, and further suggesting that the Upd3 produced by the tumours acts directly or indirectly to alter the midgut PCs.

Altogether, these different results support a model in which neoplastic tumours in *Drosophila* larvae promote cachexia-like phenotypes accompanied by alterations in the cellular composition of the larval midgut mediated by tumour-derived Ilp8 and Upd3 (Fig. 5).

**Figure 5.**
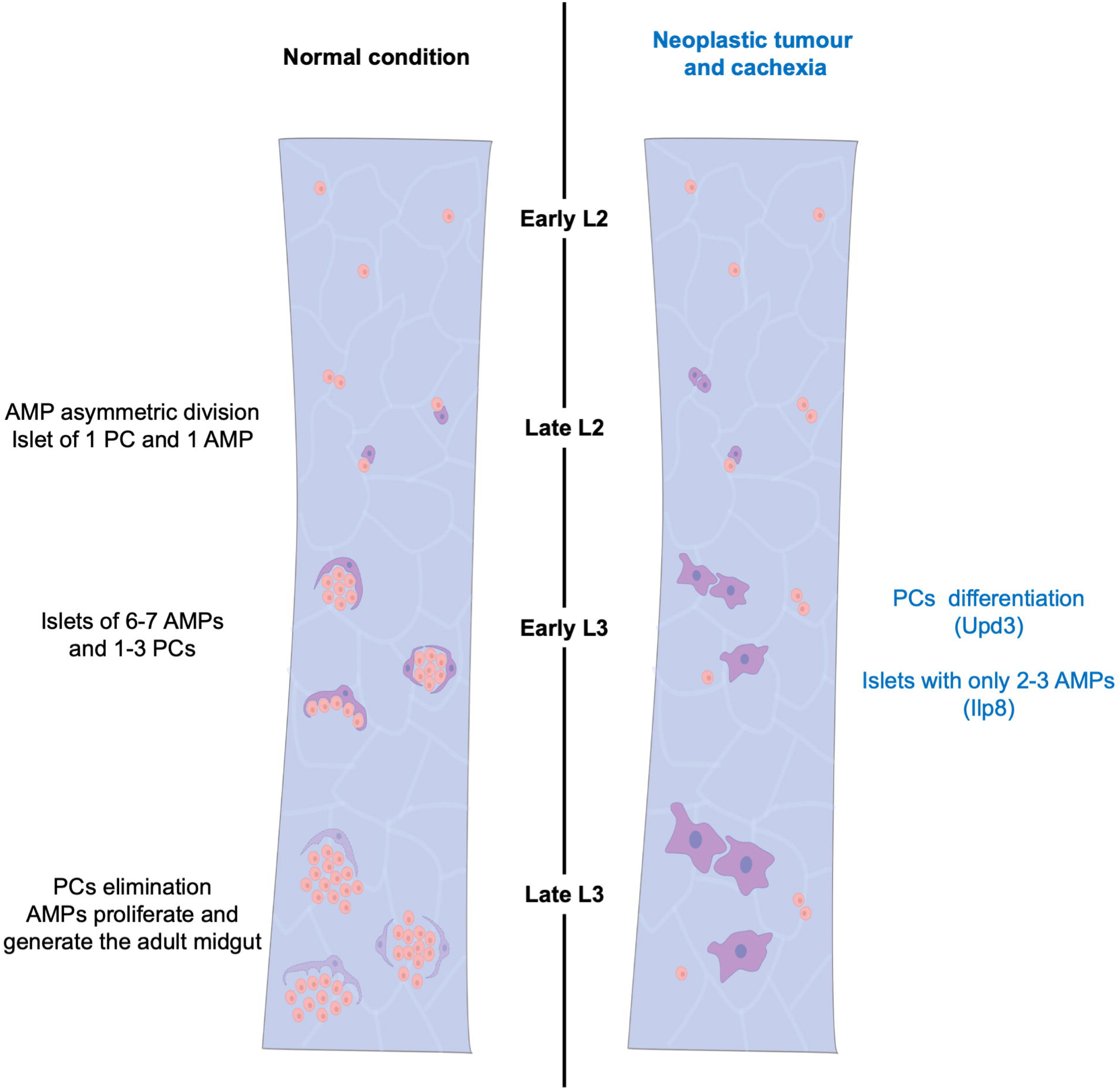
Model of tumour-associated larval midgut remodelling. The larval midgut is remodelled in response to overgrowing neoplastic wing disc tumours, through the combined action of the tumour derived secreted factors Ilp8, preventing AMP proliferation, and Upd3, promoting PC polyploidisation and differentiation.

## DISCUSSION

In this study we show that neoplastic tumours in larval wing discs based on Notch activation trigger cachexia-like syndrome in *Drosophila* larvae. These results thus extend the observations made for RasV12 eye-discs tumours (Lodge *et al*, 2021; Bakopoulos *et al*, 2023; Newton *et al*, 2020; Santabárbara-Ruiz & Léopold, 2021), and suggest that some of the features identified previously in larval models of cachexia might not be restricted to the origin of the tumour (eye disc or wing disc), or of the oncogenic driver (Ras or Notch), even though this remains to be formally tested. Indeed, Notch-based wing discs tumours do up-regulate previously identified larval cachexia mediators such as Mmp1 (Lodge *et al*, 2021) and Bnl (Newton *et al*, 2020). We further show that cachexia-inducing tumours also distantly affect the larval midgut triggering midgut cell population alterations, and in particular, a decrease in esg+ cells with a depletion of Adult Midgut Precursors (AMPs), and the polyploidization of the niche Peripheral Cells (PCs) and their engagement toward Enterocyte (EC) differentiation.

Regarding midgut cell population alterations, we show here that Ilp8, produced by the tumours, triggers a reduction in AMPs numbers per islets. AMPs clusters originate during larval life through the proliferation of initially isolated AMPs which are derived from specific posterior embryonic cells related to neuroblasts (endoblasts; (Plygawko *et al*, 2025)). Daughter cells remain together, and at each division the cluster of AMPs grows, totalling 6/7 AMPs on average in late L3. In NS cachectic larvae, we could not detect any increase in cell death in the affected midguts, suggesting that the decreased AMP numbers is unlikely the consequence of their elimination. Earlier time point analysis, revealed that AMP depletion is already present at 5d AEL. This is compatible with a model in which AMPs proliferation is slowed down leading to fewer AMPs. Supporting the decreased proliferation model, we observed a decreased expression of *polo*, coding for the polo serine/threonine kinase critical for cell proliferation, and in particular for mitosis (Pintard & Archambault, 2018). Since AMPs are the only dividing cells in the larval midgut, the polo downregulation indicates either a decrease in AMPs and/or a reduction in their mitotic potential. Interestingly, in the adult midgut, *polo* is expressed by the intestinal stem cells (ISCs), and its depletion by RNAi in ISCs promotes their depletion and differentiation into ECs (Khaminets *et al*, 2020; Zhang *et al*, 2023). Whether polo knock down in AMPs/PCs also triggers EC differentiation in the larval gut represents an attractive mechanism that could link low AMP numbers and PC polyploidization and differentiation, which remains to be formally tested. It should be noted however that PC nuclear size increase is controlled by Upd3 which does not have any impact on AMP numbers in this system, suggesting that AMP number control and PC differentiation might be two processes that could be uncoupled.

We identified Ilp8 produced by the tumours as a critical mediator of AMP numbers. Since Ilp8 is known to prevent ecdysone release in the haemolymph and thus delay developmental timing (Colombani *et al*, 2015, 2012; Garelli *et al*, 2012, 2015; Vallejo *et al*, 2015), and since ecdysone signalling is required for AMP proliferation during larval life (Micchelli *et al*, 2011), we propose that the action of tumour-derived Ilp8 on AMP numbers is indirect and involves Lgr3+ neurons in the central brain preventing ecdysone secretion by the ring gland and ecdysone signalling on AMPs. Indeed, we could observe defects in ecdysone signalling in the gut with low expression of classic ecdysone targets. Furthermore, larvae with wing disc tumours are developmentally delayed and fail to pupariate.

But the tumours also affect the larval midgut through the production of Upd3, which promotes an alteration in PCs cellular shape and nuclear size. This effect of Upd3 produced by the tumours is reminiscent to the role of Upd3 during larval midgut infection with the pathogenic *Ecc15* bacteria. There, damaged enterocytes were shown to produce Upd3 which acted locally to promote the transient and early activation of PCs and AMPs to form new enterocytes replacing the damaged ones (Houtz *et al*, 2019). Indeed, impairing *upd3* prevented AMPs depletion during infection, and overexpressing Upd3 in the AMPs and PCs (using the esg-Gal4 driver) was sufficient in normal uninfected larvae to reduce AMP numbers and drive early differentiation (Houtz *et al*, 2019). We suggest that similar mechanisms occur during tumour-associated cachexia, where the source of Upd3 is the tumour.

How is Upd3 produced by the tumours acting on the midgut? Is it direct or indirect? Previous reports showing that Jak/Stat signalling is required in *esg* positive cells for their differentiation to ECs (Houtz *et al*, 2019), supports a model where Upd3 acts on PCs.

In patients suffering from cachexia, intestine dysfunctions have been reported, and in particular gut barrier dysfunction and altered microbiome (Ni *et al*, 2021; Ubachs *et al*, 2021; Bindels *et al*, 2018). Interestingly, in the C26 mouse model of cancer-associated cachexia, a decrease in the expression of the intestinal stem cell and renewal markers Lgr5 and Klf4 was observed, suggesting that during cachexia, similarly to what we report here in *Drosophila*, cachectic tumours promote intestinal cell population alterations. The C26 model produces large amounts of IL-6, the mouse homologue of Upd3, and treating mice with an anti-IL6 antibody mitigated the weight loss and the gut permeability, but authors did not monitor whether blocking IL-6 would also prevent changes in cell population markers such as Lgr5 and Klf4 (Bindels *et al*, 2018). The critical role of tumour-derived IL-6 for cachexia in the C26 model has recently been challenged. Indeed, knocking-out *IL6* by Crispr in C26 cells delayed cachexia, in part due to increased tumour immune infiltrate slowing down the growth of the C26 grafted cells (Kwon & Hui, 2024). But, IL-6 produced by the tumours, or by other affected organs such as fat or muscles (Rupert *et al*, 2021), remains an attractive cachexia mediator and impairing IL-6 signalling, either with blocking antibodies or knocking out its receptor can mitigate cachexia in mice models (Sun *et al*, 2024; Rupert *et al*, 2021; Strassmann *et al*, 1992; White *et al*, 2011; Bindels *et al*, 2018). Importantly, in these different studies, the impact on the gut cell populations of IL-6 manipulations were not formally addressed, and further studies are thus required to investigate the impact of cachexia-inducing tumours on the host’s gut cell populations, and to assess whether gut cell population alterations represent an additional paraneoplastic symptom associated with cachexia.

## MATERIALS AND METHODS

### Drosophila genetics

All crosses were cultured at 25°C on standard food, except for the experiments with “blue-dyed” food supplemented with 1.5% of the vital blue dye Erioglaucine (Sigma Aldricht #861146). Neoplastic overgrowth of the larval imaginal discs were obtained with different models.

The first model consisted of random clones generated in the wing discs as previously described in (Djiane *et al*, 2013; Logeay *et al*, 2022) by crossing *abxUbxFLPase; Act>y>Gal4, UAS GFP; FRT82B tubGal80* flies with flies carrying either *FRT82B* (Ctrl), or *UAS-Nicd; FRT82B scrib1* (Ubx>NS neoplastic discs).

The second model consisted of driving by *Sd-Gal4* (wing pouch) *UAS-Nicd* and the *scrib* RNAi *P{TRiP.HMS01490}attP2* (Sd>NS neoplastic discs).

Information on gene models and functions, and on *Drosophila* lines available were obtained from FlyBase (flybase.org – (Thurmond *et al*, 2019)). Stocks used are listed below:

Bloomington Drosophila Stock Center: BDSC

Vienna Drosophila Resource Center: VDRC

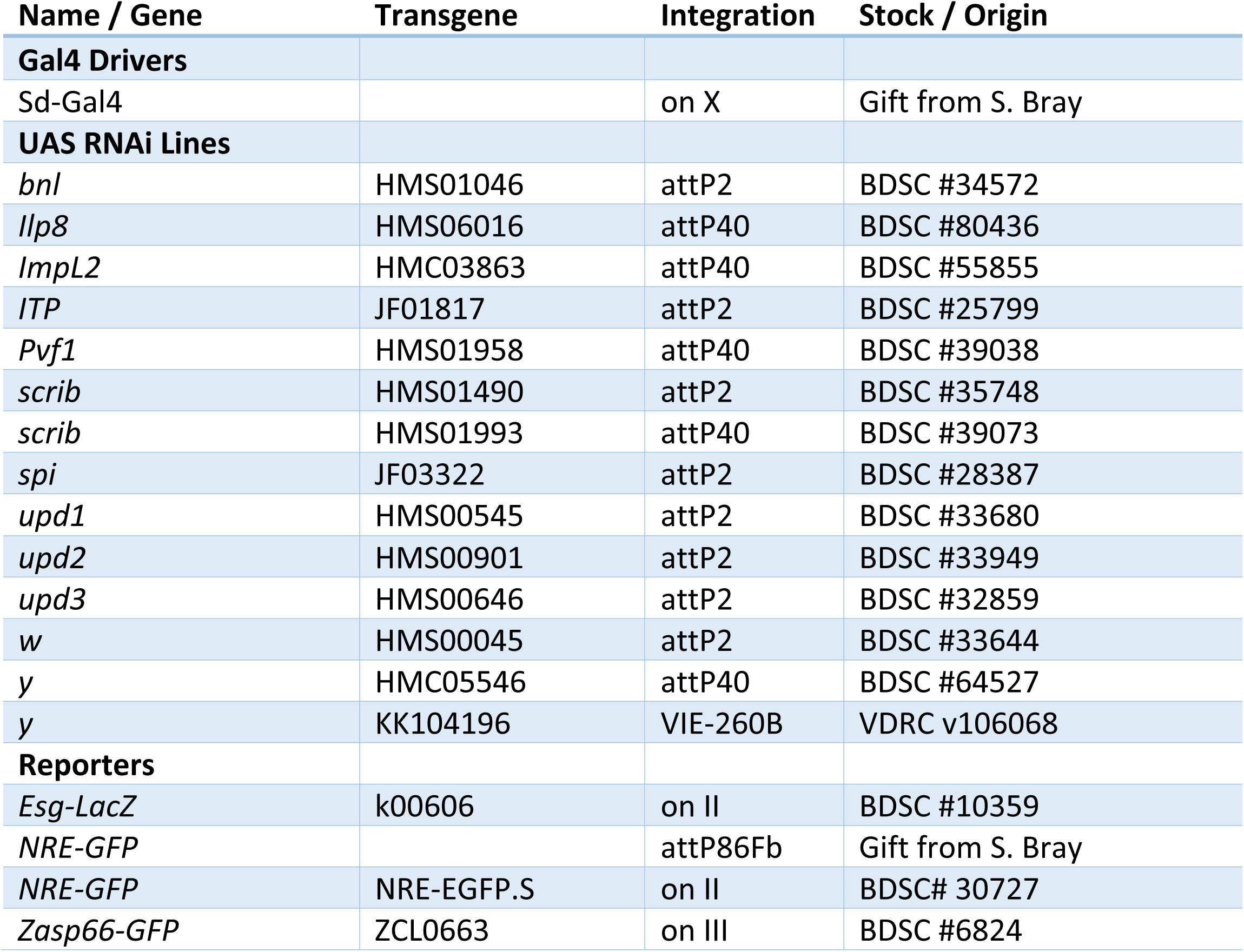

### Haemolymph extraction and trehalose measurement

3rd instar larvae were washed in water, in ethanol 70% EtOH, and dried. Larvae were placed on Parafilm on ice and pierced carefully with forceps. Haemolymph was collected by pipetting. To remove debris, the haemolymph was centrifuged for 30min at 1000g at 4C and the supernatant was further centrifuged at 4C for 20min at 15000g. 1.5µl of haemolymph was used to measure trehalose according to (Strassburger & Teleman, 2016) and using the Glucose assay kit (Sigma-Aldrich #GAGO20).

### Glycogen and glucose measurements in fat body

Dissected fat bodies pooled from 10 3rd instar larvae were homogenized for glycogen and glucose measurements according to (Strassburger & Teleman, 2016) and using the Glucose assay kit (Sigma-Aldrich #GAGO20).

### Immunofluorescence

Tissues were dissected in PBS and fixed in 4% formaldehyde for 30min (discs) or 4-12h (midguts) at room temperature (RT). They were washed 3x 10min in PBT (PBS 0,2% Triton X-100), and blocked in PBT-BSA (PBT 5% BSA) for 30min. Primary antibodies were incubated overnight at 4C in PBT-BSA. Tissues were washed 3x 10min in PBT, and secondary antibodies were incubated 2h at RT in PBT-BSA. They were washed 3×10 min in PBT and mounted in CitiFluor^TM^ AF1 (Agar). Images were acquired on an upright Leica THUNDER or a Zeiss Apotome microscope.

Primary antibodies were: Mouse anti-armadillo (1:200; Developmental Study Hybridoma Bank – DSHB #N2 7A1), Rat anti-ECadherin (1:25; DSHB #DCAD2), Rabbit anti-GFP (1:500; Torrey Pines Biolabs #TP401), Mouse anti-b-galactosidase (1:200; DSHB #40-1a), Rabbit anti-b-galactosidase (1:200; MPBIO 085597-CF), Mouse anti-prospero (1:500; DSHB #MR1A), Rabbit anti-Cleaved Dcp-1 (1:500; Cell Signaling #9578). Secondary antibodies conjugated to Alexa Fluor 488, 555, 647, or to Cy3 were from Jackson Labs Immuno Research (1:200). DAPI was used at 1:2000 and Phalloidin-Rhodamine (Sigma-Aldrich #P1951) was used at 1:200.

### Quantifications

#### AMP numbers without Esg staining

Within the first region of the midgut, small cells arranged in islet, not positively stained for Prospero were counted using the Cell Counter plugging in FIJI (ImageJ). Numbers were corrected by the area of the gut.

#### PC engulfment and detachment

Within the first region of the midgut, number of misshapen and normal-looking PC were counted. PC were considered misshapen and isolated if they did not project their membrane around AMPs. Percentage of isolated from total number of PC was calculated.

### Nuclear size

The nuclei of NRE-GFP positive cells were captured and measured on FIJI (ImageJ)

#### Lipid droplets staining with BODIPY

Fat bodies from 3^rd^ instar larvae were dissected in PBS and fixed in 4% formaldehyde for 30min at RT. Tissues were then rinsed 2x in PBS and incubated in BODIPY FL Dye (1:500 dilution from a 1mg/mL stock, ThermoFisher) for 30min in PBS. They were rinsed 2x in PBS and immediately mounted in CitiFluor^TM^ AF1 (Agar). Images were acquired on a Leica THUNDER microscope. Lipid droplet numbers and sizes were measured using FIJI (ImageJ) software.

#### Western blot

Western blot analyses were performed according to standard protocols. Primary antibodies used in this study: Mouse anti-α-Actinin (1:2000; DSHB #2G3-3D7), Mouse anti-Myosin (1:2000; DSHB #3E8-3D3), Rabbit anti-Lipophorin II (1:15000; gift from J Culi). Secondary antibodies conjugated to HRP were from Jackson Labs Immuno Research (1:15000). For WB on haemolymph, 5µl of haemolymph with 5µl of 2X Laemmli were deposited per lane.

#### Quantitative RT-PCR

For each dissected tissue, pooled groups of ten were lysed chemically with QIAzol Lysis Reagent (QIAGEN) and physically using a tissue disruptor Retsch MM400 (Verder Scientific). Total RNA was then purified using QIAGEN kits (QIAGEN RNeasy Lipid #74804 or RNeasy Plus #74134). Genomic DNA was removed by incubating with DNase (QIAGEN #79254), and cDNA was retro-transcribed using SuperScript III (Invitrogen #18080-044). Semi quantitative RT-qPCR was performed on biological triplicates using SYBR Green I Master mix on a LightCycler^®^ 480 (Roche). Fold change was estimated using the ΔΔCT approach.

Primers used:

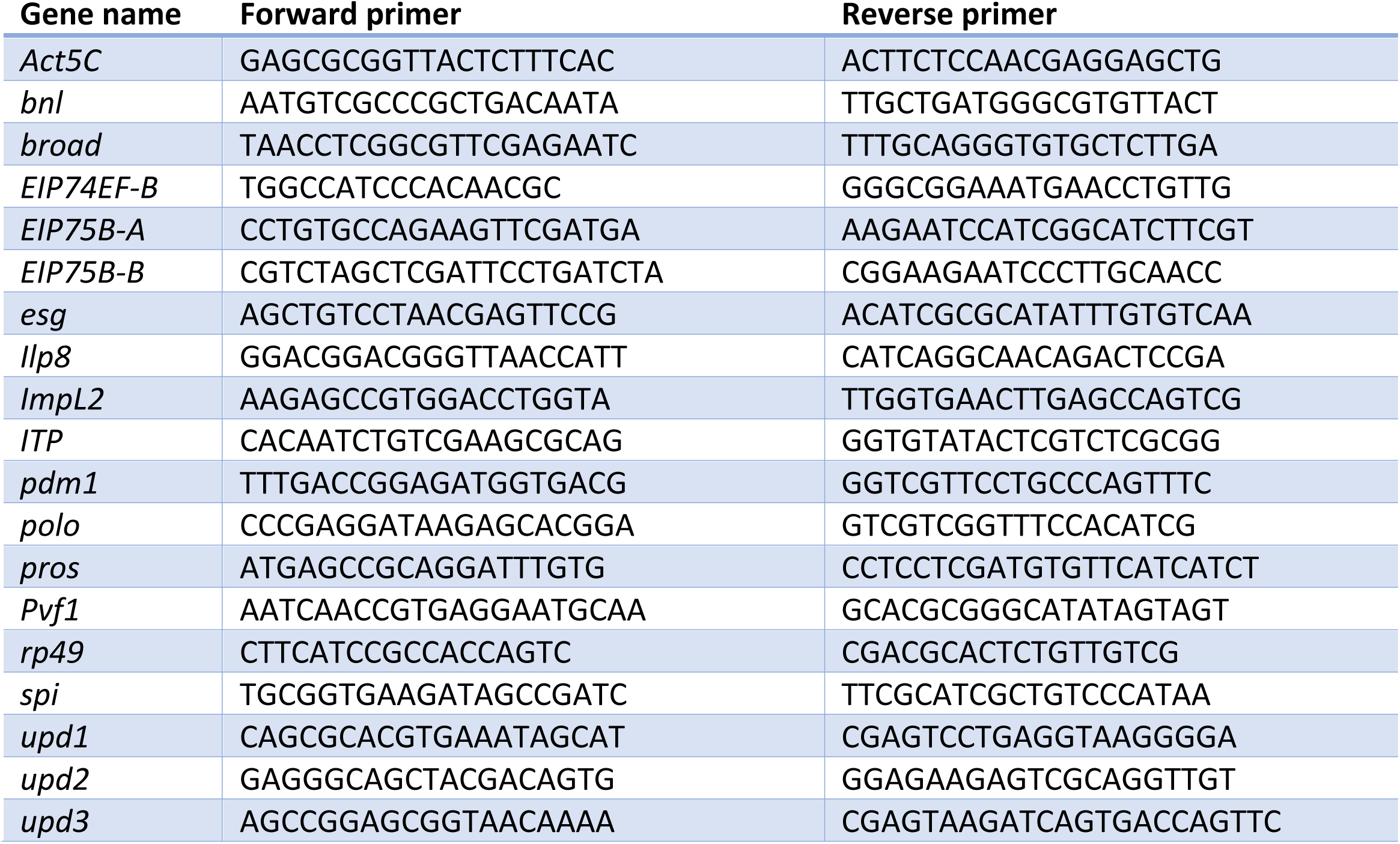

## ACKNOWLEDGEMENTS

We thank K. Basler, S. Bray, J. Culi, M. Milán, and P. Léopold for sharing flies, and A. Gallet and J. Pannequin for insightful discussions We acknowledge the Bloomington Drosophila Stock Center (BDSC - NIH P40OD018537), the Vienna Drosophila Resource Center (VDRC), the Developmental Studies Hybridoma Bank, the Montpellier Drosophila facility, the Montpellier Imaging facility (MRI), and FlyBase for providing reagents and tools critical for our research.

## AUTHOR CONTRIBUTIONS

Conceptualization: AD. Methodology: CG, AD. Validation: JF, MRV, KS, CG. Formal analysis: JF, CG, AD. Investigation: JF, MRV, KS, CJT, LHM, PL, CG, AD. Writing – original Draft: AD. Writing – review and editing: JF, CG, AD. Visualization: JF, CG, AD. Supervision: CG, AD. Project administration: AD. Funding acquisition: CG, AD.

## DATA AVAILABILITY

RNA-Seq data have been deposited in NCBI’s Gene Expression Omnibus under accession number GSE185339.

---

**Supplemental Figure S1.**
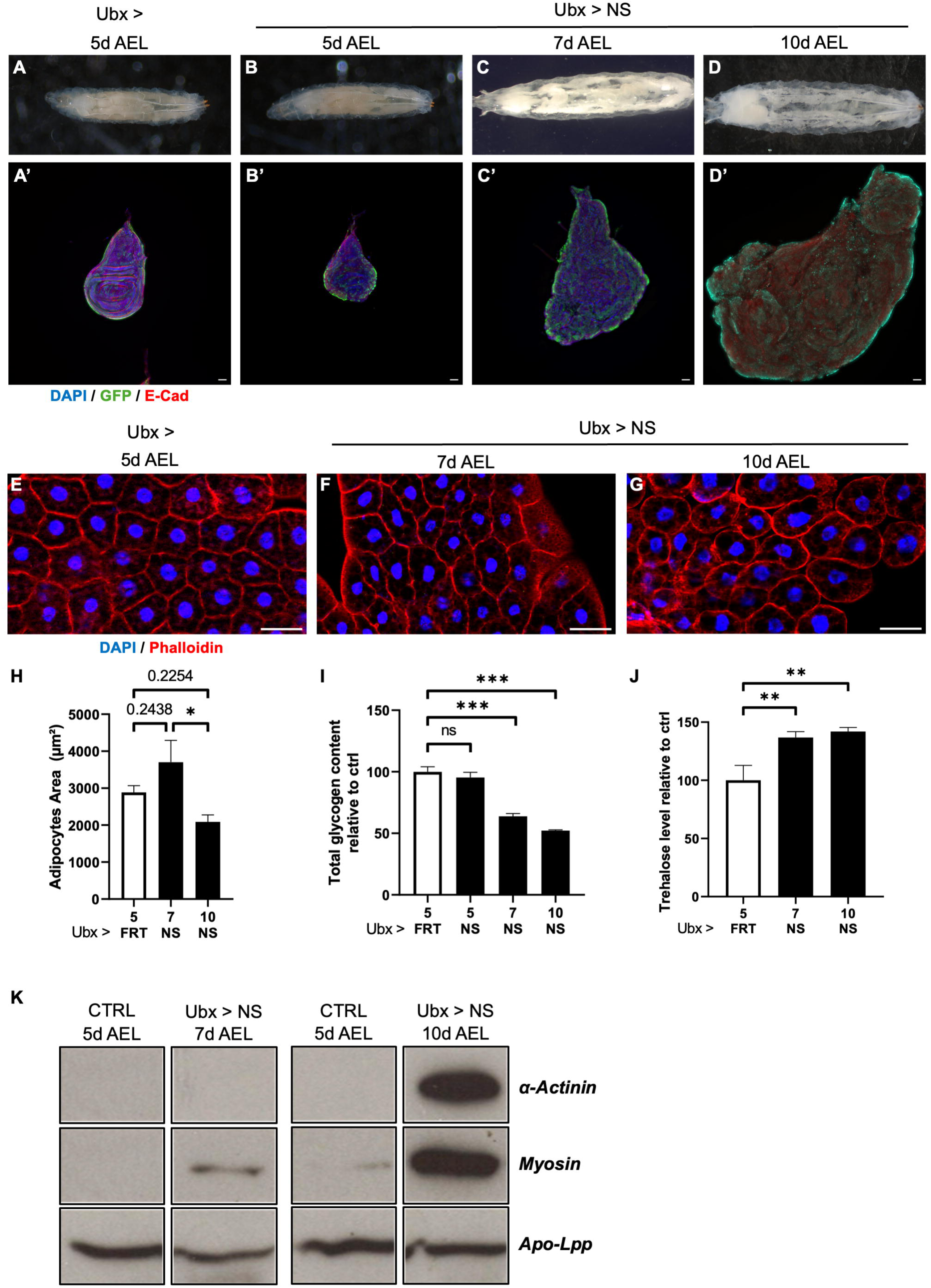
UbxMARCM NS larvae exhibit cachexia-like syndrome. **A-D.** Timeline of larvae morphology in response to neoplastic wing disc tumours with activated Notch and mutant for scribble (Ubx>NS) at 5d (B), 7d (C), and 10d AEL (D) compared to wild type 5d AEL Ubx> controls (A). (A-D) Whole larva, head on the left. (A’-D’) Corresponding dissected wing imaginal discs and stained for GFP (clones of over-expression, green), E-Cadherin (membranes, E-Cad, red), and DAPI (nuclei, blue). White bar: 50 µm. **E-G.** Staining of larval fat body showing Actin (red), and nuclei (DAPI, blue). Control Ubx> larvae at 5d AEL (E), and cachectic Ubx>NS larvae at 7d AEL (F) and 10d AEL (G). White bar: 50 µm. **H.** Quantification of adipocyte cell size from images shown in (E-G). Biological replicates n ≥ 4. Error-bars show standard error of the mean (sem). Normality was tested with a Shapiro-Wilk Test. One-way ANOVA statistical test. * p < 0.05. **I.** Quantification of glycogen content in whole larvae (mainly in fat body). Biological replicates n = 6. Error-bars show standard error of the mean (sem). Normality was tested with a Shapiro-Wilk Test. One-way ANOVA statistical test. *** p < 0.001, ns = not significant. **J.** Quantification of trehalose concentration in the haemolymph. Biological replicates n = 3. Error-bars show standard error of the mean (sem). One-way ANOVA statistical test. *** p < 0.001, ** p < 0.01. **K.** Western blot of whole protein extracts of hemolymph (5µl samples) from Ubx> control larvae at 5d AEL, and Ubx>NS cachectic larvae at 7d and 10d AEL, and monitoring the presence of muscle proteins such as Myosin and α-Actinin. Apo-Lpp is used as loading control.

**Supplemental Figure S2.**
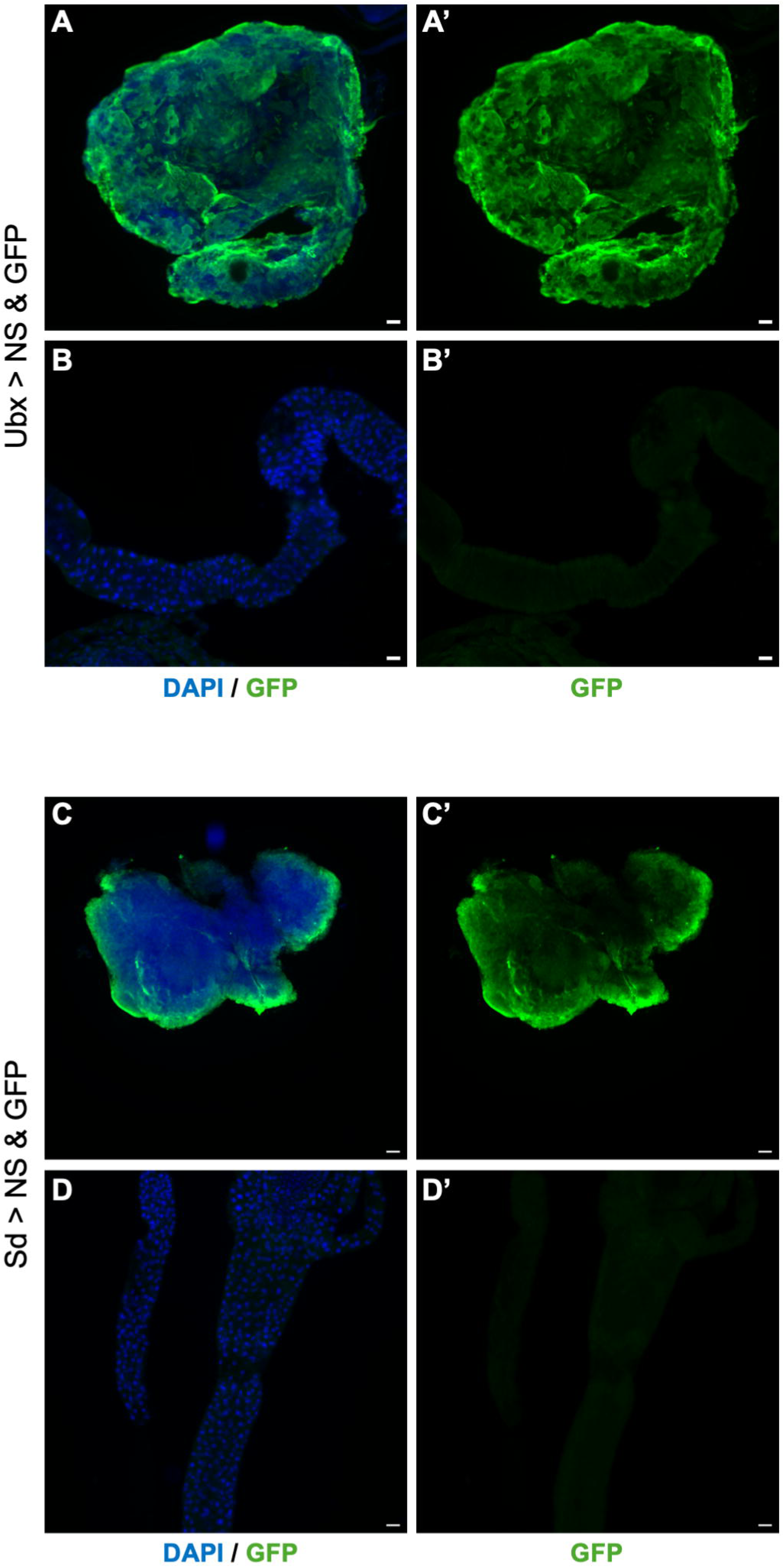
UbxMARCM and Sd drivers are not expressed in the larval midgut. **A-B.** GFP driven in the Ubx>NS model and showing dissected wing disc (A) and midgut (B). GFP (green in A-B and in A’-B’), DAPI (blue in A-B). White bar: 50 µm in A, and 100µm in B. **C-D.** GFP driven in the Sd>NS model and showing dissected wing disc (C) and midgut (D). GFP (green in C-D and in C’-D’), DAPI (blue in C-D). White bar: 50 µm in C, and 100µm in D.

**Supplemental Figure S3.**
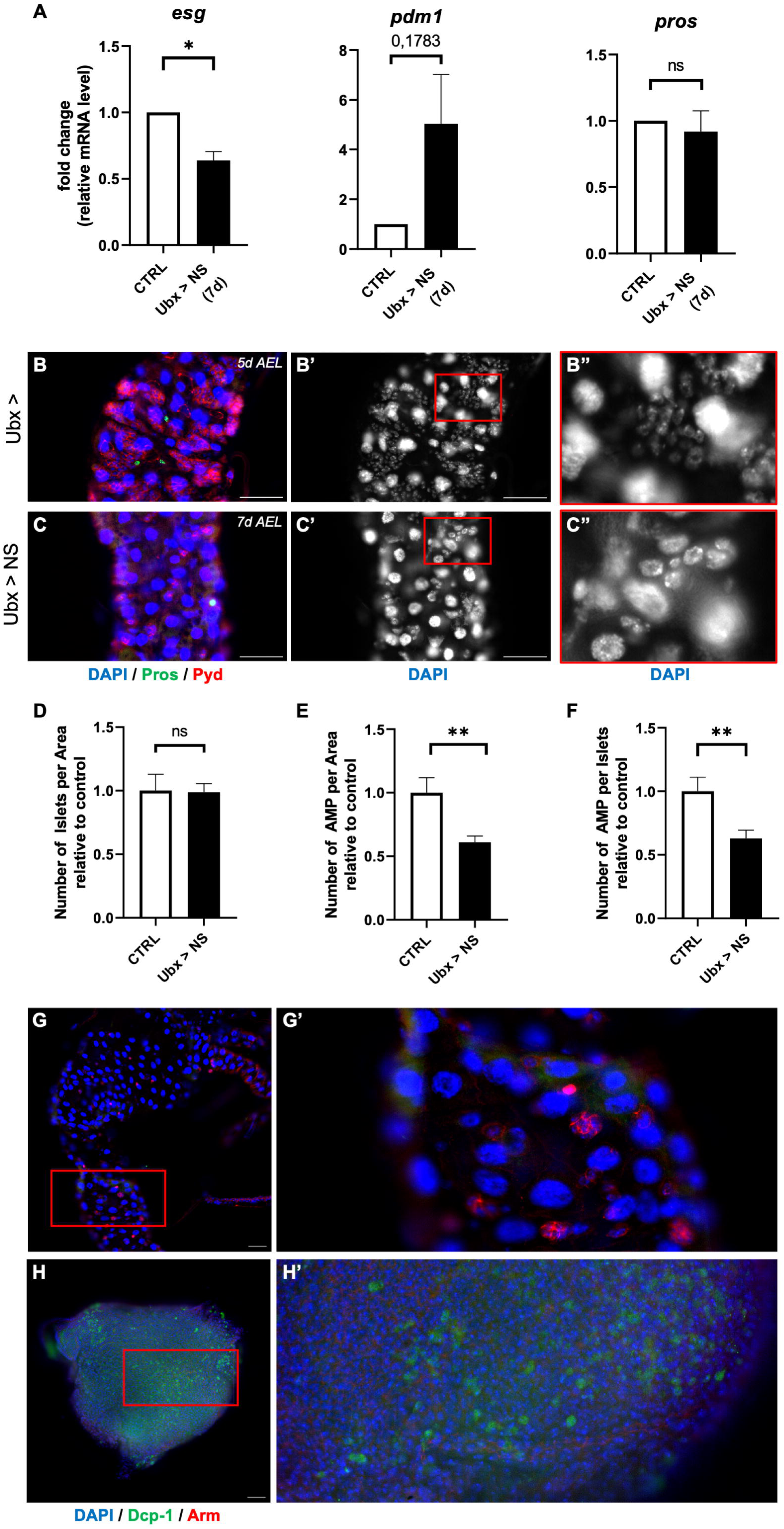
Low AMP numbers in the midguts of Ubx>NS cachectic larvae. **A.** Expression of the cell type markers *esg*, *pdm1*, and *pros* in Ubx>NS larval midguts at 7d AEL and monitored by qRT-PCR. Reference gene for normalization: *rp49/RpL32*. Biological replicates n = 3. Error-bars show standard error of the mean (sem). Normality was tested with a Shapiro-Wilk Test. One-way ANOVA statistical test. * p < 0.05, ns = not significant. **B-C.** Staining of larval midguts at 7d AEL showing the membrane marker Armadillo (red), and nuclei (DAPI, blue in B-C, white in B’-C’ and in B’’-C’’); white bar: 50 µm. (B’’-C’’) higher magnification of images in (B’-C’); AMPs are characterised by clusters of cells with small nuclei. **D-F.** Quantification of number of AMP islets per area unit (D), of AMP per area unit (E), and of AMP per islet (F) in Ubx>NS midguts from images shown in (B-C). Biological replicates n = 3. Error-bars show standard error of the mean (sem). One-way ANOVA statistical test. ** p < 0.01, ns = not significant. **G-H.** Activated Dcp-1 caspase staining in larval midgut (G) and wing imaginal discs (H) in Sd>NS cachectic larvae at 7d AEL. Midguts and wing discs were stained for Dcp-1 (activated caspase, green), the membrane marker Armadillo (red), and nuclei (DAPI, blue). (G’-H’) higher magnification of images in (G-H). White bar: 100 µm in G, 50 µm in H.

**Supplemental Figure S4.**
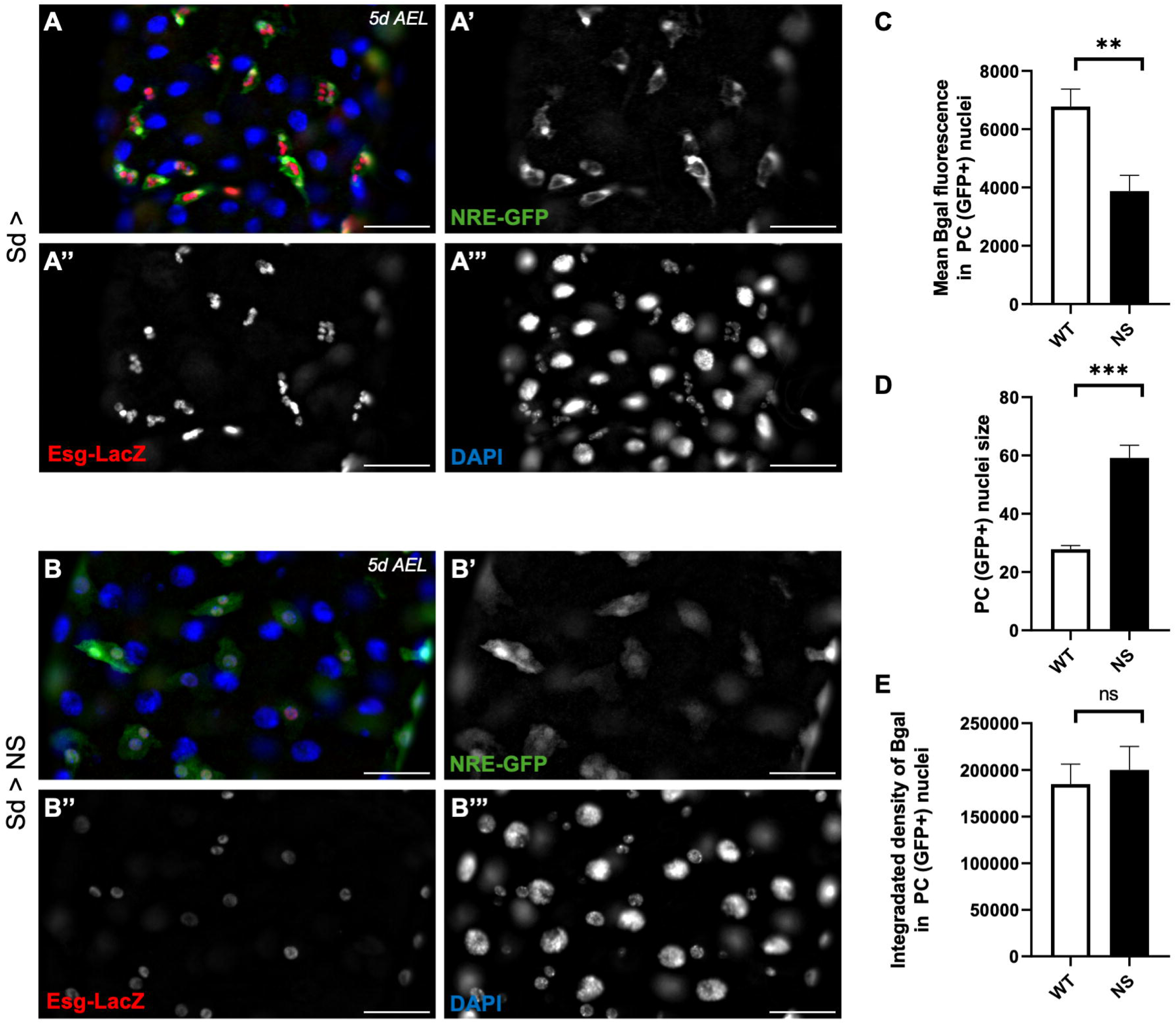
Esg-LacZ expression in misshapen PCs. **A-B.** Larval midgut staining showing the PC marker NRE-GFP (green in A-B, white in A’-B’), the AMP&PC marker Esg (Esg-LacZ, red in A-B, white in A’’-B’’), and nuclei (DAPI, blue in A-B, white in A’’’-B’’’). Control Sd-Gal4 midguts (A) and cachectic Sd>NS midguts (B) at 5d AEL. White bar: 50 µm. **C.** Quantification of mean nuclear β-galactosidase fluorescence intensity in PCs from control and NS guts. T-test statistical test ; **p < 0.01. **D.** Quantification of nuclear size in PCs from control and NS guts. Normality was tested with a Shapiro-Wilk Test. T-test statistical test ; ***p < 0.001. **E.** Integrated density of nuclear β-galactosidase signal showing no significant difference between control and NS PCs. Normality was tested with a Shapiro-Wilk Test. T-test statistical test ; ns = not significant.

**Supplemental Figure S5.**
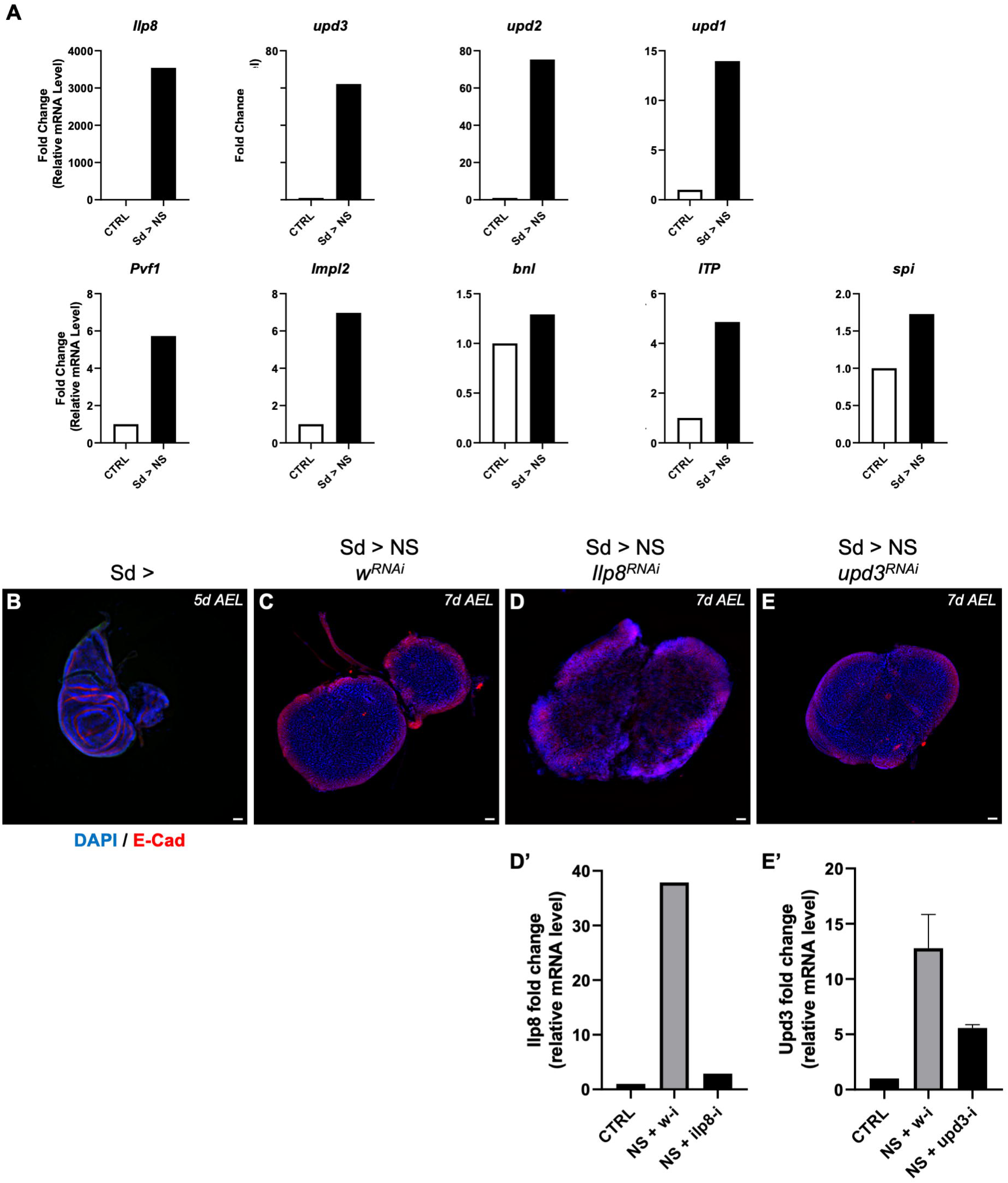
Cachectic tumours derived secreted factors. **A.** Validation in Sd>NS neoplastic wing discs of the up-regulation of the secreted factors identified in the RNA-Seq transcriptome of Ubx>NS wing discs. Expression was monitored by qRT-PCR compared to control Sd> wing discs. Reference gene for normalization: *rp49/RpL32*. **B-E.** Staining of Sd> control wing discs (B), and Sd>NS neoplastic wing discs at 7d AEL expressing either a control white RNAi (C), an RNAi targeting *Ilp8* (D), or an RNAi targeting *upd3* (E). Discs were stained for for Actin (red), and DAPI (nuclei, blue). White bar: 50 µm. (D’-E’). Evaluation of the knock-down efficiency of the *Ilp8* (D’) and *upd3* (E’) RNAi used monitored by qRT-PCR. Reference gene for normalization: *rp49/RpL32*. Error-bars show standard error of the mean (sem).

## REFERENCES

Baazim H, Antonio-Herrera L & Bergthaler A (2022) The interplay of immunology and cachexia in infection and cancer. Nat Rev Immunol 22: 309–321

Bakopoulos D, Golenkina S, Dark C, Christie EL, Sánchez-Sánchez BJ, Stramer BM & Cheng LY (2023) Convergent insulin and TGF-β signalling drives cancer cachexia by promoting aberrant fat body ECM accumulation in a Drosophila tumour model. EMBO Rep 24: e57695

Baracos VE, Martin L, Korc M, Guttridge DC & Fearon KCH (2018) Cancer-associated cachexia. Nat Rev Dis Primers 4: 17105

Beltrà M, Pöllänen N, Fornelli C, Tondla K, Hsu MY, Zampieri S, Moletta L, Corrà S, Porporato PE, Kivelä R, et al (2023) NAD+ repletion with niacin counteracts cancer cachexia. Nat Commun 14: 1849

Bilgic SN, Domaniku A, Toledo B, Agca S, Weber BZC, Arabaci DH, Ozornek Z, Lause P, Thissen J-P, Loumaye A, et al (2023) EDA2R-NIK signalling promotes muscle atrophy linked to cancer cachexia. Nature 617: 827–834

Bindels LB, Neyrinck AM, Loumaye A, Catry E, Walgrave H, Cherbuy C, Leclercq S, Van Hul M, Plovier H, Pachikian B, et al (2018) Increased gut permeability in cancer cachexia: mechanisms and clinical relevance. Oncotarget 9: 18224–18238

Boumard B & Bardin AJ (2021) An amuse-bouche of stem cell regulation: Underlying principles and mechanisms from adult Drosophila intestinal stem cells. Curr Opin Cell Biol 73: 58–68

Burfeind KG, Zhu X, Norgard MA, Levasseur PR, Huisman C, Buenafe AC, Olson B, Michaelis KA, Torres ER, Jeng S, et al (2020) Circulating myeloid cells invade the central nervous system to mediate cachexia during pancreatic cancer. Elife 9: e54095

Colombani J, Andersen DS, Boulan L, Boone E, Romero N, Virolle V, Texada M & Léopold P (2015) Drosophila Lgr3 Couples Organ Growth with Maturation and Ensures Developmental Stability. Curr Biol 25: 2723–2729

Colombani J, Andersen DS & Léopold P (2012) Secreted peptide Dilp8 coordinates Drosophila tissue growth with developmental timing. Science 336: 582–585

Ding G, Xiang X, Hu Y, Xiao G, Chen Y, Binari R, Comjean A, Li J, Rushworth E, Fu Z, et al (2021) Coordination of tumor growth and host wasting by tumor-derived Upd3. Cell Rep 36: 109553

Djiane A, Krejci A, Bernard F, Fexova S, Millen K & Bray SJ (2013) Dissecting the mechanisms of Notch induced hyperplasia. EMBO J 32: 60–71

Ferrer M, Anthony TG, Ayres JS, Biffi G, Brown JC, Caan BJ, Cespedes Feliciano EM, Coll AP, Dunne RF, Goncalves MD, et al (2023a) Cachexia: A systemic consequence of progressive, unresolved disease. Cell 186: 1824–1845

Ferrer M, Mourikis N, Davidson EE, Kleeman SO, Zaccaria M, Habel J, Rubino R, Gao Q, Flint TR, Young L, et al (2023b) Ketogenic diet promotes tumor ferroptosis but induces relative corticosterone deficiency that accelerates cachexia. Cell Metab 35: 1147–1162.e7

Figueroa-Clarevega A & Bilder D (2015) Malignant Drosophila tumors interrupt insulin signaling to induce cachexia-like wasting. Dev Cell 33: 47–55

Garelli A, Gontijo AM, Miguela V, Caparros E & Dominguez M (2012) Imaginal discs secrete insulin-like peptide 8 to mediate plasticity of growth and maturation. Science 336: 579–582

Garelli A, Heredia F, Casimiro AP, Macedo A, Nunes C, Garcez M, Dias ARM, Volonte YA, Uhlmann T, Caparros E, et al (2015) Dilp8 requires the neuronal relaxin receptor Lgr3 to couple growth to developmental timing. Nat Commun 6: 8732

Hodgson JA, Parvy J-P, Yu Y, Vidal M & Cordero JB (2021) Drosophila Larval Models of Invasive Tumorigenesis for In Vivo Studies on Tumour/Peripheral Host Tissue Interactions during Cancer Cachexia. Int J Mol Sci 22: 8317

Holland P, Quintana EM, Khezri R, Schoborg TA & Rusten TE (2022) Computed tomography with segmentation and quantification of individual organs in a D. melanogaster tumor model. Sci Rep 12: 2056

Houtz P, Bonfini A, Bing X & Buchon N (2019) Recruitment of Adult Precursor Cells Underlies Limited Repair of the Infected Larval Midgut in Drosophila. Cell Host Microbe 26: 412–425.e5

Khaminets A, Ronnen-Oron T, Baldauf M, Meier E & Jasper H (2020) Cohesin controls intestinal stem cell identity by maintaining association of Escargot with target promoters. Elife 9: e48160

Khezri R, Holland P, Schoborg TA, Abramovich I, Takáts S, Dillard C, Jain A, O’Farrell F, Schultz SW, Hagopian WM, et al (2021) Host autophagy mediates organ wasting and nutrient mobilization for tumor growth. EMBO J 40: e107336

Kir S, White JP, Kleiner S, Kazak L, Cohen P, Baracos VE & Spiegelman BM (2014) Tumour-derived PTH-related protein triggers adipose tissue browning and cancer cachexia. Nature 513: 100–104

Kwok SH, Liu Y, Bilder D & Kim J (2024) Paraneoplastic renal dysfunction in fly cancer models driven by inflammatory activation of stem cells. Proc Natl Acad Sci U S A 121: e2405860121

Kwon Y, Song W, Droujinine IA, Hu Y, Asara JM & Perrimon N (2015) Systemic organ wasting induced by localized expression of the secreted insulin/IGF antagonist ImpL2. Dev Cell 33: 36–46

Kwon Y-Y & Hui S (2024) IL-6 promotes tumor growth through immune evasion but is dispensable for cachexia. EMBO Rep 25: 2592–2609

Lemaitre B & Miguel-Aliaga I (2013) The digestive tract of Drosophila melanogaster. Annu Rev Genet 47: 377–404

Liu X, Li S, Cui Q, Guo B, Ding W, Liu J, Quan L, Li X, Xie P, Jin L, et al (2024) Activation of GPR81 by lactate drives tumour-induced cachexia. Nat Metab 6: 708–723

Lodge W, Zavortink M, Golenkina S, Froldi F, Dark C, Cheung S, Parker BL, Blazev R, Bakopoulos D, Christie EL, et al (2021) Tumor-derived MMPs regulate cachexia in a Drosophila cancer model. Dev Cell 56: 2664–2680.e6

Logeay R, Géminard C, Lassus P, Rodríguez-Vázquez M, Kantar D, Heron-Milhavet L, Fischer B, Bray SJ, Colinge J & Djiane A (2022) Mechanisms underlying the cooperation between loss of epithelial polarity and Notch signaling during neoplastic growth in Drosophila. Development 149: dev200110

Mathur D, Bost A, Driver I & Ohlstein B (2010) A transient niche regulates the specification of Drosophila intestinal stem cells. Science 327: 210–213

Micchelli CA (2012) The origin of intestinal stem cells in Drosophila. Dev Dyn 241: 85–91

Micchelli CA & Perrimon N (2006) Evidence that stem cells reside in the adult Drosophila midgut epithelium. Nature 439: 475–479

Micchelli CA, Sudmeier L, Perrimon N, Tang S & Beehler-Evans R (2011) Identification of adult midgut precursors in Drosophila. Gene Expr PaQerns 11: 12–21

Miguel-Aliaga I, Jasper H & Lemaitre B (2018) Anatomy and Physiology of the Digestive Tract of Drosophila melanogaster. GeneRcs 210: 357–396

Newton H, Wang Y-F, Camplese L, Mokochinski JB, Kramer HB, Brown AEX, Fets L & Hirabayashi S (2020) Systemic muscle wasting and coordinated tumour response drive tumourigenesis. Nat Commun 11: 4653

Ni Y, Lohinai Z, Heshiki Y, Dome B, Moldvay J, Dulka E, Galffy G, Berta J, Weiss GJ, Sommer MOA, et al (2021) Distinct composition and metabolic functions of human gut microbiota are associated with cachexia in lung cancer patients. ISME J 15: 3207– 3220

Ohlstein B & Spradling A (2006) The adult Drosophila posterior midgut is maintained by pluripotent stem cells. Nature 439: 470–474

Overend G, Luo Y, Henderson L, Douglas AE, Davies SA & Dow JAT (2016) Molecular mechanism and functional significance of acid generation in the Drosophila midgut. Sci Rep 6: 27242

Pintard L & Archambault V (2018) A unified view of spatio-temporal control of mitotic entry: Polo kinase as the key. Open Biol 8: 180114

Plygawko AT, Stephan-Otto Attolini C, Pitsidianaki I, Cook DP, Darby AC & Campbell K (2025) The Drosophila adult midgut progenitor cells arise from asymmetric divisions of neuroblast-like cells. Dev Cell 60: 429–446.e6

Rodríguez-Vázquez M, Falconi J, Heron-Milhavet L, Lassus P, Géminard C & Djiane A (2024) Fat body glycolysis defects inhibit mTOR and promote distant muscle disorganization through TNF-α/egr and ImpL2 signaling in Drosophila larvae. EMBO Rep 25: 4410– 4432

Romão D, Muzzopappa M, Barrio L & Milán M (2021) The Upd3 cytokine couples inflammation to maturation defects in Drosophila. Curr Biol 31: 1780–1787.e6

Rupert JE, Narasimhan A, Jengelley DHA, Jiang Y, Liu J, Au E, Silverman LM, Sandusky G, Bonetto A, Cao S, et al (2021) Tumor-derived IL-6 and trans-signaling among tumor, fat, and muscle mediate pancreatic cancer cachexia. J Exp Med 218: e20190450

Saavedra P, Dumesic PA, Hu Y, Filine E, Jouandin P, Binari R, Wilensky SE, Rodiger J, Wang H, Chen W, et al (2023) REPTOR and CREBRF encode key regulators of muscle energy metabolism. Nat Commun 14: 4943

Saavedra P & Perrimon N (2019) Drosophila as a Model for Tumor-Induced Organ Wasting. Adv Exp Med Biol 1167: 191–205

Santabárbara-Ruiz P & Léopold P (2021) An Oatp transporter-mediated steroid sink promotes tumor-induced cachexia in Drosophila. Dev Cell 56: 2741–2751.e7

Song W, Kir S, Hong S, Hu Y, Wang X, Binari R, Tang H-W, Chung V, Banks AS, Spiegelman B, et al (2019) Tumor-Derived Ligands Trigger Tumor Growth and Host Wasting via Differential MEK AcAvaAon. Dev Cell 48: 277–286.e6

Strassburger K & Teleman AA (2016) Protocols to Study Growth and Metabolism in Drosophila. Methods Mol Biol 1478: 279–290

Strassmann G, Fong M, Kenney JS & Jacob CO (1992) Evidence for the involvement of interleukin 6 in experimental cancer cachexia. J Clin Invest 89: 1681–1684

Sun Q, van de Lisdonk D, Ferrer M, Gegenhuber B, Wu M, Park Y, Tuveson DA, Tollkuhn J, Janowitz T & Li B (2024) Area postrema neurons mediate interleukin-6 function in cancer cachexia. Nat Commun 15: 4682

Thurmond J, Goodman JL, Strelets VB, Attrill H, Gramates LS, Marygold SJ, Matthews BB, Millburn G, Antonazzo G, Trovisco V, et al (2019) FlyBase 2.0: the next generation. Nucleic Acids Res 47: D759–D765

Ubachs J, Ziemons J, Soons Z, Aarnoutse R, van Dijk DPJ, Penders J, van Helvoort A, Smidt ML, Kruitwagen RFPM, Baade-Corpelijn L, et al (2021) Gut microbiota and short-chain fatty acid alterations in cachectic cancer patients. J Cachexia Sarcopenia Muscle 12: 2007–2021

Vallejo DM, Juarez-Carreño S, Bolivar J, Morante J & Dominguez M (2015) A brain circuit that synchronizes growth and maturation revealed through Dilp8 binding to Lgr3. Science 350: aac6767

White JP, Baynes JW, Welle SL, Kostek MC, Matesic LE, Sato S & Carson JA (2011) The regulation of skeletal muscle protein turnover during the progression of cancer cachexia in the Apc(Min/+) mouse. PLoS One 6: e24650

Xie H, Heier C, Meng X, Bakiri L, Pototschnig I, Tang Z, Schauer S, Baumgartner VJ, Grabner GF, Schabbauer G, et al (2022) An immune-sympathetic neuron communication axis guides adipose tissue browning in cancer-associated cachexia. Proc Natl Acad Sci U S A 119: e2112840119

Xu J, Liu Y, Yang F, Cao Y, Chen W, Li JSS, Zhang S, Comjean A, Hu Y & Perrimon N (2024) Mechanistic characterization of a Drosophila model of paraneoplastic nephrotic syndrome. Nat Commun 15: 1241

Xu W, Li G, Chen Y, Ye X & Song W (2023) A novel antidiuretic hormone governs tumour-induced renal dysfunction. Nature 624: 425–432

Yeom E, Shin H, Yoo W, Jun E, Kim S, Hong SH, Kwon D-W, Ryu TH, Suh JM, Kim SC, et al (2021) Tumour-derived Dilp8/INSL3 induces cancer anorexia by regulating feeding neuropeptides via Lgr3/8 in the brain. Nat Cell Biol 23: 172–183

Zhang Y, Chen R, Gong L, Huang W, Li P, Zhai Z & Ling E (2023) Regulation of intestinal stem cell activity by a mitotic cell cycle regulator Polo in Drosophila. G3 (Bethesda) 13: jkad084

Zhou X, Wang JL, Lu J, Song Y, Kwak KS, Jiao Q, Rosenfeld R, Chen Q, Boone T, Simonet WS, et al (2010) Reversal of cancer cachexia and muscle wasting by ActRIIB antagonism leads to prolonged survival. Cell 142: 531–543

